# Auditory training alters the cortical representation of complex sounds

**DOI:** 10.1101/2023.12.29.573645

**Authors:** Huriye Atilgan, Kerry M Walker, Andrew J. King, Jan W. Schnupp, Jennifer K. Bizley

## Abstract

Auditory learning is supported by long-term changes in the neural processing of sound. We examined these task-depend changes in auditory cortex by mapping neural sensitivity to timbre, pitch and location cues in trained ferrets (n = 5), and untrained control ferrets (n = 5). Trained animals either identified vowels in a two-alternative forced choice task (n = 3) or discriminated when a repeating vowel changed in identity or pitch (n = 2). Neural responses were recorded under anesthesia in two primary auditory cortical fields and two tonotopically organized non-primary fields. In trained animals, the overall sensitivity to sound timbre was reduced across three cortical fields compared to control animals, but maintained in a non-primary field (the posterior pseudosylvian field). While training did not increase sensitivity to timbre across auditory cortex, it did change the way in which neurons integrated spectral information with neural responses in trained animals increasing their sensitivity to first and second formant frequencies,, whereas in control animals’ cortical sensitivity to spectral timbre depends mostly on the second formant. Animals trained on timbre identification were required to generalize across pitch when discriminating timbre and their neurons became less modulated by fundamental frequency relative to control animals. Finally, both trained groups showed increased spatial sensitivity and an enhanced response to sound source locations close to the midline, where the loudspeaker was located in the training chamber. These results demonstrate that training elicited widespread alterations in the cortical representation of complex sounds.

## Introduction

Sensory discrimination tasks are known to drive cortical plasticity, and increases in the representational area of the stimulus, such as tonotopic map expansion, have been proposed as providing the structural substrate for learning in the auditory cortex (Rutkowski and Weinberger, 2005; Polley et al., 2006; Engineer et al., 2014; Schreiner and Polley, 2014). However, other studies have noted that such changes only occur for simple stimulus features such as frequency or level, and questioned the functional role of this form of representational plasticity; plasticity may be a temporary phenomenon associated with learning that does not persist once a task is well learned (Reed et al., 2011), and frequency discrimination can occur in the absence of map plasticity (Brown, 2004).

Learning to discriminate behaviourally meaningful sounds, such as when mothers learn to recognize pup vocalisations, also elicits changes in auditory cortical neurons, altering inhibition and leading to gain enhancement independently of any changes in tonotopic representation. Animals trained to dynamically locate a target area based on properties of sound lead to adaptive changes within the primary auditory cortex (A1) (Bao et al., 2004; Whitton et al., 2014). For example, locating a weak tone in the presence of background noise led to increased noise tolerance, and an increase in the number of non-monotonic rate-level functions (Whitton et al., 2014) while navigating based on the repetition rate of brief sound bursts led to changes in temporal response properties (Bao et al., 2004).

Studies investigating training induced changes after learning to make discriminations based on complex sound features (i.e. for sounds composed of multiple sound frequencies) have reported changes in the way that neurons optimize integration of both spectral and temporal features. For example, training cats to discriminate changes in the position of spectral peaks in a harmonic sound complex (akin to single formant vowels) led to neurons sharpening their frequency tuning and shifting their spectrotemporal preference towards trained sounds, suggestive of altered spectral integration (Keeling et al., 2008). Training ferrets to discriminate forward from reversed vocalisations increased the information conveyed in auditory cortical temporal pattern codes in trained relative to naïve animals (Schnupp et al., 2006). In non-human primates trained to detect increases in the rate of sound amplitude modulation, auditory cortical neurons showed narrower spectral tuning and a shift in preference towards the trained modulation rates (Beitel et al., 2020).

While most studies of representational plasticity have focused on A1, studies investigating short-term plasticity during active behaviour reveal receptive field plasticity that is more marked in higher cortical areas compared to primary auditory cortex (Atiani et al., 2014; Elgueda et al., 2019). Moreover, higher cortical areas are more likely to show selectivity to spectrally complex sounds such as speech (Mohn et al., 2024)This raises the question of whether longer-term changes in the way in which auditory cortical neurons represent trained stimuli might also vary across the auditory cortical hierarchy.

In this study, we recorded from the auditory cortex of adult ferrets (n = 5), trained to discriminate the timbre of spectrally overlapping artificial vowels, and compared neural responses to the same sounds recorded in naïve animals (n = 5). Animals were trained to identify vowel timbre in the context of a two-alternative forced choice (n = 3) task (Bizley et al., 2013; Town et al., 2018) or discriminate timbre or pitch in a go/no-go task (n = 2, Walker et al., 2017). After behavioral training was complete, electrophysiological recordings were made from four tonotopic auditory cortical fields. Previous work shows that the perceptual features of complex sounds, such as their location in space or spectral timbre, are distributed across auditory cortical fields (Bizley et al., 2009; Walker et al., 2011; Allen et al., 2017). Wesought to determine: (i) how training altered single neuron response sensitivity to both learned and passively exposed sound features; and (ii) whether training induced effects differed across the auditory heirachy. Our data demonstrate that training alters the way in which neurons integrate spectral information, and the neural representations of trained and passively exposed features are affected in different ways. Specifically, while timbre sensitivity was maintained in a non-primary field (the posterior pseudosylvian field, PPF) in trained animals, neurons in other fields became markedly less sensitive to timbre. In contrast, sensitivity to sound location, which was an untrained but task-relevant feature, was enhanced in trained animals.

## Materials & Methods

### Animals

All animal procedures were approved by the Committee on Animal Welfare and Ethical Review at the University of Oxford and performed under license from the UK Home Office in accordance with the Animal (Scientific Procedures) Act 1986. Ten adult, female, pigmented ferrets (*Mustela putorius*) were used in this study. Five of these were naïve control animals, from whom some data was previously published (Bizley et al., 2009; Walker et al., 2011). Five of these were trained animals, three of the experimental animals experienced 1-2 years of training on a two-alternative forced-choice timbre identification task, which required them to report the identity of an artificial vowel (Fig. 1C, T-Id). For extended details of behavioral training see “stimuli” (below), (Walker et al., 2009), and Figure 1. Two of the experimental animals were trained to perform a go/no-go task, in which they were presented with a sequence of artificial vowels and had to discriminate either changes in vowel identity, or in different sessions, changes in vowel F0 (Fig.1D, TP-Disc). For more details on this training paradigm, see “stimuli” (below) and Figure 1 (Walker et al., 2011).

**Figure 1:**
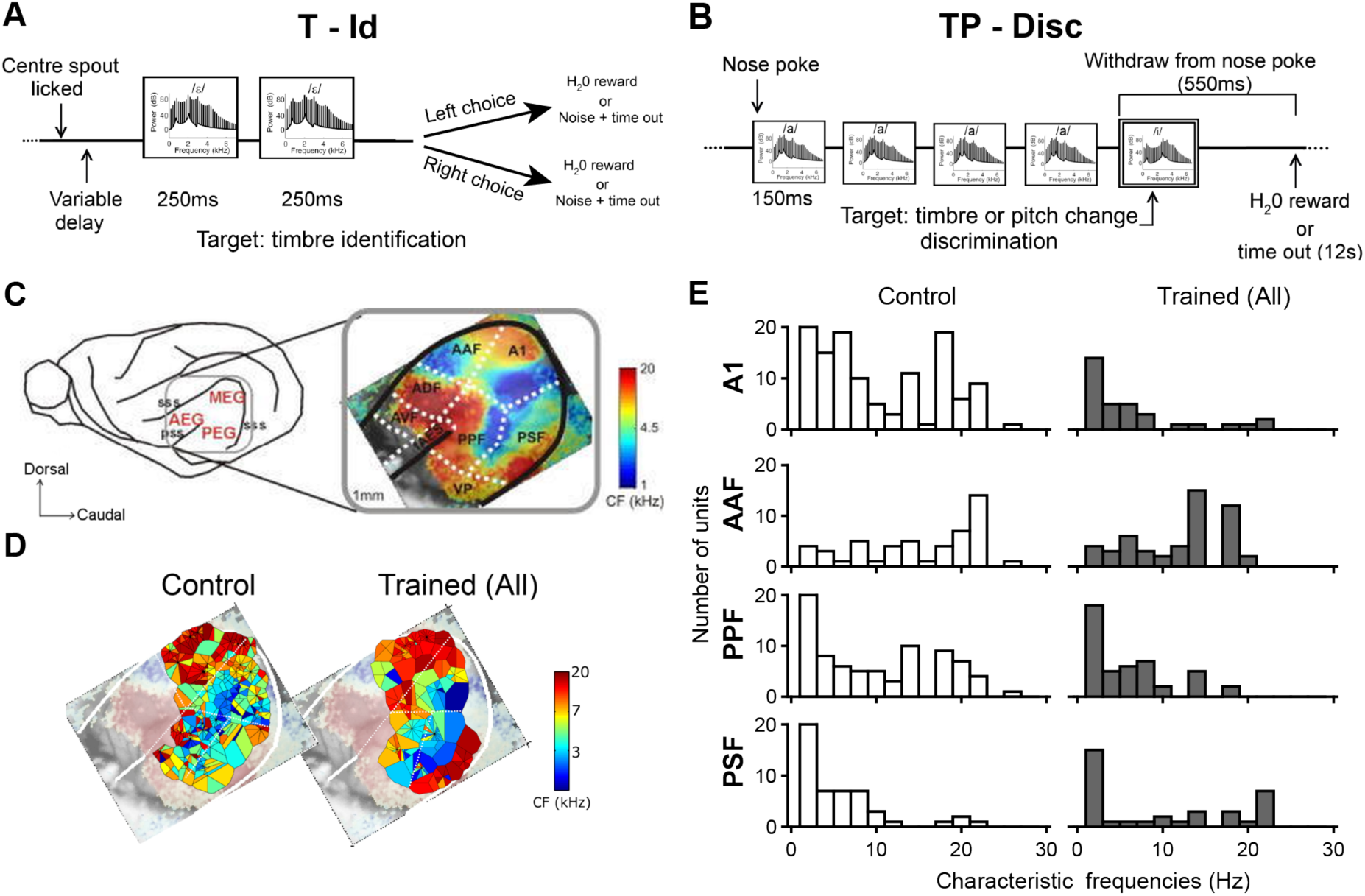
Tonotopic organization of auditory cortex of ferret and behavioral paradigm. **A-B** Schematic illustration of the timbre two-alternative forced choice paradigm (A, T-Id) and timbre/pitch discrimination paradigm (B, TP-Disc). **C**, Location of ferret auditory cortical fields and their tonotopic organization (adapted from (Nelken et al., 2004). Recordings in this study targeted primary auditory cortex (A1), anterior auditory field (AAF), posterior pseudosylvian field (PPF), and posterior suprasylvian field (PSF). Field boundaries are marked with dotted lines, and the pseudosylvian sulcus (pss) and suprasylvian sulcus (sss) are drawn as solid lines. **D**, Voronoi tessellation map showing the CFs of all unit recordings made in control animals (615 units, 5 animals) and trained animals (783 units, 5 animals). Tiles represent a recording site and are colored according to the characteristic frequency (CF) of the unit recorded there. **E**, Histograms of the characteristic frequency distribution across subregions of the auditory cortex: A1, AAF, PPF and PSF. Data from control animals are on the left, and data from all trained animals are on the right. The histograms represent the number of units responsive to specific frequency ranges (Hz) within each cortical area.

Ferrets were housed in groups of either two or three, with free access to high-protein food pellets and water bottles. For experimental ferrets, on the day before behavioral training, water bottles were removed from the home cages and were replaced on the last afternoon of a training run. Training lasted for 5 days or less, with at least 2 days between each run. On training days, ferrets received drinking water as positive reinforcement while performing a sound discrimination task. Water consumption during training was measured and supplemented as wet food in home cages at the end of the day to ensure that each ferret received at least 60 ml of water per kilogram of body weight daily. Once behavioral training was complete, electrophysiological recordings were made under non-recovery anesthesia (details below). Recording under anesthesia was necessary for the large-scale mapping of neurons across cortical fields, and in order to directly compare the resulting responses with data from control animals previously collected under the same anesthetic regime.

### Stimuli

Acoustic stimuli for both behavioral testing and electrophysiology were artificial vowel sounds. For electrophysiological testing, sounds were all possible combinations of four F0 values (F0 = 200, 336, 565, and 951 Hz), four spectral timbres (/a/: F1–F4 at 936, 1551, 2815, and 4290 Hz; /ε/: 730, 2058, 2979, and 4294 Hz; /u/: 460, 1105, 2735, and 4115 Hz; and /i/: 437, 2761, 3372, and 4352 Hz), and four locations presented in virtual acoustic space (-45°, - 15°, 15°, and 45° azimuth, all at 0° elevation). This gave a total of 64 sounds, each of which was 150 ms in duration. Although the animals were trained using different stimulus protocols, recordings were conducted for all stimulus combinations in both the T-Id and TP-Disc groups, which included all vowels used in both training regimens. Additionally, noise bursts and pure tones were used to characterize individual units, and to determine tonotopic gradients in order to confirm the cortical field in which any given recording was made (Bizley et al., 2005).

### Behavioral Testing

Full details of the training apparatus and procedure for shaping animals are provided in Walker et al. (2011) and Bizley et al. (2013). Briefly, in the timbre identification (T-Id) task water-restricted ferrets were positively conditioned in a two-alternative forced choice task to report the identity of a vowel sound, which could be either /u/ (F1-F4 460, 1105, 2735, 4155 Hz, animals conditioned to respond at the left spout) or /ε/: (F1-F4: 730, 2058, 2979, 4294, animals conditioned to respond at the right spout). Animals were trained initially with an F0 of 200 Hz, but tested across a range of values from 150 Hz – 500 Hz.

In the timbre/pitch discrimination (TP-Disc) task, water-restricted ferrets were trained to report a change in the pitch or timbre of a repeating artificial vowel in a go/no-go task . The reference sound was the vowel /a/ (F1–F4: 936, 1551, 2815, 4290 Hz) with a F0 of 200 Hz, and F0 targets were the vowel /a/ with F0 values of 336, 565 and 951 Hz, while timbre targets were the vowels /i/, /u/ and /ε/, presented at an F0 of 200 Hz.

In both tasks, the animal initiated each trial by inserting its nose in a poke hole situated at the center of the sound-isolated testing chamber. For T-Id task, this resulted in the presentation of two identical repetitions of one of the vowel sounds, and animals were rewarded for correctly responding at the side that was associated with that vowel. In the TP-Disc task, ferrets heard a sequence of artificial vowels, which could change in identity or pitch at the third to seventh vowel in the sequence, and if ferrets withdrew from the nose poke hole during the presentation of such a deviant, they were rewarded with water. Failures to withdraw to a deviant (within a 550 ms time window following deviant onset) resulted in a 12-s time out. In both tasks, sounds were presented from a loudspeaker located 5 cm above the central ‘go’ spout at the animals’ midline (0° azimuth).

### Electrophysiological Recordings

Experimental methods were identical to those reported in Bizley et al. (2009),which comprises the control dataset for this study. Anesthesia was induced by a single dose of a mixture of medetomidine (Domitor; 0.022 mg/kg/h; Pfizer) and ketamine (Ketaset; 5 mg/kg/h; Fort Dodge Animal Health). The left radial vein was cannulated and a continuous infusion (5 ml/h) of a mixture of medetomidine and ketamine in physiological saline containing 5% glucose was provided throughout the experiment. The ferrets also received a single, subcutaneous, dose of 0.06 mg/kg/h atropine sulfate (C-Vet Veterinary Products, UK) to reduce bronchial secretions and, every 12 h, subcutaneous doses of 0.5 mg/kg dexamethasone (Dexadreson; Intervet UK) to reduce cerebral oedema. The ferret was intubated, placed on a ventilator (7025 respirator; Ugo Basile, Italy) and supplemented with oxygen. Body temperature, end-tidal CO_2_, and the electrocardiogram (ECG) were monitored throughout the experiment. Electrophysiology experiments typically lasted between 36 and 60 h.

The animal was placed in a stereotaxic frame and the temporal muscles on both sides were retracted to expose the dorsal and lateral parts of the skull. A metal bar was cemented and screwed into the right side of the skull, holding the head without further need of a stereotaxic frame. On the left side, the temporal muscle was largely removed, and the suprasylvian and pseudosylvian sulci were exposed by a craniotomy, exposing auditory cortex. The dura was retracted and the cortex covered with silicon oil or agar. The animal was then transferred to a small table in an anechoic chamber (IAC, UK). Sounds presented through customized Panasonic RPHV297 headphone drivers. Closed-field calibrations were performed using a one-eighth inch condenser microphone (Brüel and Kjær), placed at the end of a model ferret ear canal, to create an inverse filter that ensured the driver produced a flat (<±5 dB) output.

Recordings targeted primary and non-primary tonotopic areas: primary auditory cortex (A1) and the anterior auditory field (AAF) on the middle ectosylvian gyrus, and the posterior pseudosylvian and posterior suprasylvian fields (PPF and PSF) located on the posterior ectosylvian gyrus (Fig. 1A). Recordings were made with silicon probe electrodes (Neuronexus Technologies, USA) either in a 16 x 2 configuration (16 active sites spaced at 100μm intervals on each of two shanks), a 32 × 1 configuration (a single shank with 50 μm spacing between sites) or, in one animal, an 8 × 4 configuration (100μm spacing in depth, 200 μm between shanks). All probes had active sites of 177μm. Voltage signals were bandpass filtered (500- 5000 Hz), amplified, and digitized at 25 kHz. Data acquisition and stimulus generation were performed using BrainWare (Tucker-Davis Technologies). Spike sorting was performed via WavClus (Quiroga et al., 2004) to isolate single units (based on waveform shape and refractory period) and multiunit clusters; we refer to ‘units’ to include both unless otherwise specified. A minimum of 80 units were recorded in each field, in each group (mean = 118 units ± 28.32, Table 1).

**Table 1:**
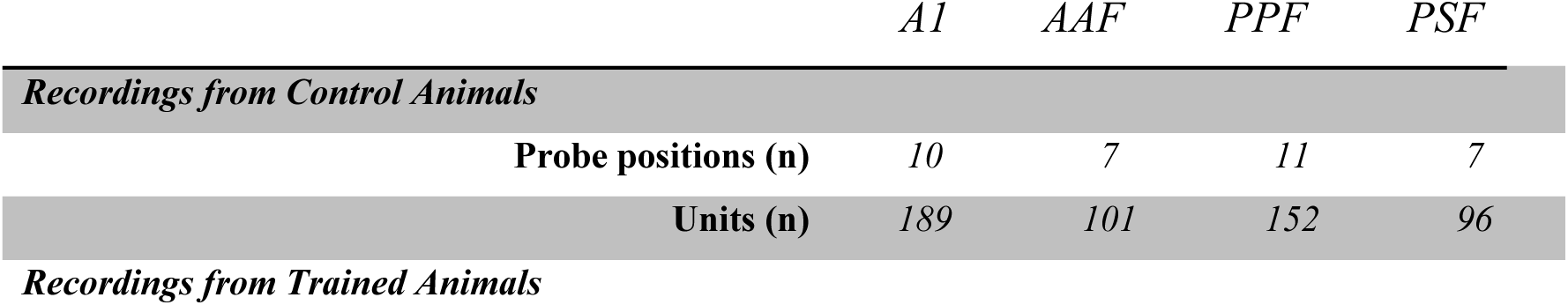
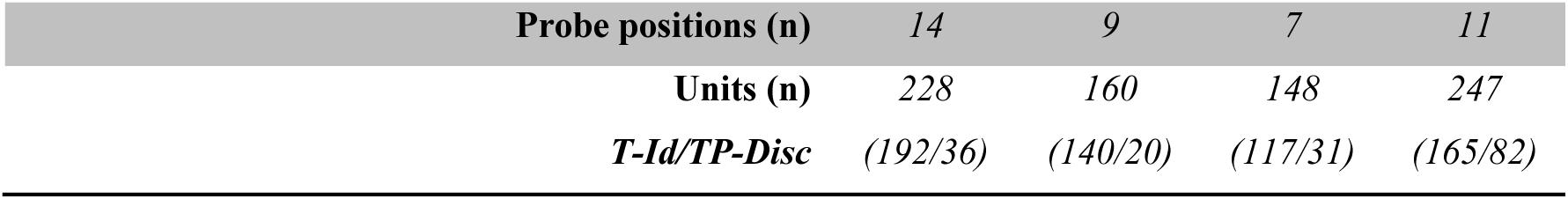
Total number of recordings (probe positions and units) in each field for 5 control animals and 5 trained animals.

Data were combined across animals within each group (control or trained) to make composite tonotopic maps (Fig. 1B). The Characteristic Frequency (the frequency to which the unit responded at its threshold sound level, CF) of each unit was determined based on its response to pure tones. CF information, along side latencies and photographs of the electrode penetration positions on the brain, to make a composite tonotopic map based on the known tonotopic organization of ferret auditory cortex (Bizley et al., 2005; Bimbard et al., 2018).

### Neural Data Analysis

Cortical responses were analyzed using a variance decomposition approach developed in Bizley et al., 2009. We first calculated spike counts for each of the 64 stimuli, averaged over repeated presentations of the same sound, and binned with 20 ms resolution over the 300 ms immediately preceding stimulus onset. We then performed a 4-way ANOVA on the spike counts, where the 3 stimulus parameters (timbre, pitch and location) plus the time bin served as factors. To quantify the relative strength with which one of the three stimulus dimensions influenced the firing of a particular unit, we calculated the proportion of variance explained by each of timbre, pitch, and location, 𝑉𝑎𝑟*_stim_*, as:

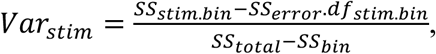

where “stim” refers to the stimulus parameter of interest (timbre, pitch and location), *SS_stim.bin_* is the sum of squares for the interaction of the stimulus parameter and time bin, *SS_error_* is the sum of squares of the error term, *df_stim.bin_* refers to the degrees of freedom for the stimulus × time bin interaction, *SS_total_* is the total sum of squares, and *SS_bin_* is the sum of squares for the time bin factor. A large *SS_bin_* reflects the fact that the response rate was not flat over the duration of the 300 ms response window. By examining the stimulus-by-time-bin interactions, we were able to test the statistical significance of the influence a given stimulus parameter had on the temporal discharge pattern of the response. Subtracting the *SS*_error_. *df_stim.bin_* from the *SS_stim.bin_* term allows us to calculate the proportion of response variance attributable to each of the stimuli, taking into account the additional variance explained simply by adding extra parameters to the model. As in our previous work (Bizley et al., 2009), we considered a main effect or interaction term in the ANOVA to be statistically significant if *p* < 0.001.

### Statistical analysis

For statistical comparison of neural sensitivity measures between groups and cortical fields (derived using the variance decomposition approach described above) we performed Generalized Mixed Linear Model regression, with the specific analysis for each test reported in the Results section. We utilized the *lmerTest* package in R. defining the model formula as ’Value ∼ Training Group * Field + (1|Penetration)’. Here, ’Value’ represented the dependent variable, while ’Training Group’ and ’Field’ were the independent variables (fixed effects). ’Penetration’ was included as random effects in the model, to take into account that simultaneous recordings from a single recording electrode are unlikely to be fully statistically independent. The model was fit using the lmer function from the *lmerTest* package. To evaluate how robustly our model accounted for the data we used five-fold cross validation, calculating the root mean square error, which indicates the model’s prediction error, for each test set.

To analyse how acoustic features shaped sensitivity to timbre across groups and cortical fields, we extended our mixed effects model to incorporate additional fixed effects (the difference in first formant frequencies between vowel in a pair, denoted ΔF1, and the difference in second formant frequencies between each vowel in a pair and interaction terms, denoted ΔF2). The updated model formula was defined as ’Value ∼ Field + Training Group + ΔF1 + ΔF2 + Training Group: ΔF1 + Training Group: ΔF2 + Field: ΔF1 + Field: ΔF2 + ΔF1: ΔF2 + (1|Unit) + (1|Penetration)’. In this expanded model, ’ΔF1’ and ’ΔF2’ represent additional fixed effects, and interactions between ’Training Group’, ’Field’, ’ΔF1’, and ’ΔF2’ were included to examine potential synergistic or antagonistic effects among these variables. We included both ’Unit’ (reflecting the multiple observations per unit) and ’Penetration’ as random effects and 5-fold cross validation was used to calculate the goodness-of-fit of the model.

To generate predictions for arbitrary ΔF1 and ΔF2 values, we used the model to predict responses to a dense grid of formant frequency space. Specifically, a range of ΔF1 values was defined by 200 equally spaced points between the minimum and maximum observed ΔF1 values in the dataset, while the range of Δ2 values comprised 400 equally spaced points within its observed range. This grid spanned the complete acoustic space represented in our dataset, allowing for a comprehensive prediction of the model’s response to changes in formant frequencies. For each unique combination of cortical field and training group, the model predictions were computed across the entire grid of F1 and F2 values, resulting in a detailed landscape of predicted timbre sensitivity. The *predict* function within *lmerTest* was employed to estimate the timbre sensitivity for each observation in the generated dataset, based on the fitted mixed-effects model. This expansive prediction dataset formed the basis for subsequent visualization and interpretation, aiming to elucidate the influence of auditory training on the neural encoding of vowel sounds.

To examine differences in spatial tuning that emerged during training we used a mixed effects model that considered the effect of (sound) ‘Location’, ’Training Group’, and ’Field’ on spike rate. The model formula was defined as ’Spike Rate ∼ Location * Training Group * Field +(1|Unit) + (1|Penetration)’, where ’Unit’ and ’Penetration’ were again included as random effects. In this model, ’Location’ was treated as a factor variable with four levels: "-45", "-15", "15", and "45", in which “-45” was explicitly used as reference. This final model allowed us to closely examine how changes in ’Location’ interacted with ’Training Group’ and ’Field’ to affect spike rate, while controlling for the random effects of ’Unit’ and ’Penetration’.

### Code and data availability

All code is available on Github: https://github.com/huriyeatg/trainingInducedPlasticity Data are available on the following repository [link added on publication].

## Results

Three ferrets were trained in a two-alternative forced choice timbre identification (T-Id) task to discriminate /u/ from /ε/, across a range of F0s (Fig.1A). Two additional ferrets were trained in a go/no-go task to discriminate changes in timbre or F0 (TP-Disc) within a repeating reference vowel (/a/, F0 = 200 Hz, Fig.1B). Once behavioral training and testing were complete, we recorded neural activity under medetomidine/ketamine anesthesia, which allowed us to map neural responses across the surface of multiple auditory cortical fields in each animal. These responses were directly compared to those obtained in naïve animals in a previous study, which constitutes the control data for this investigation. The activity of 783 units (459 single neurons, 324 multi-units), which were responsive to vowels (paired t-test on sound-evoked and spontaneous firing rates, *p<0.05*), were recorded from four tonotopic auditory cortical fields (see Table 1 and Fig. 1C, D), and compared to driven 538 units from control animals. The CF distribution was not significantly different between trained and naïve animals (Fig. 1E, GLM to predict CF with factors field (A1, AAF, PPF, PSF) and group (Control/TD-Disc/T-ID): and their interactions, all p values >0.27).

As in the control data presented in Bizley et al., 2009, single units displayed diverse time- varying responses to the complete stimulus set (Fig. 2, see also extended data). For instance, Unit A in Fig. 2A displayed a very selective response to the 200 Hz fundamental frequency with a high first formants (for 936 Hz and 730 Hz). In contrast, another unit (Figure 2A, Unit B) exhibited a distinctive response pattern characterized by enhanced firing rate for a high first formant onset (the highest for 936 Hz, followed by 730 Hz) and a reduced firing rate during the stimulus.

**Figure 2.**
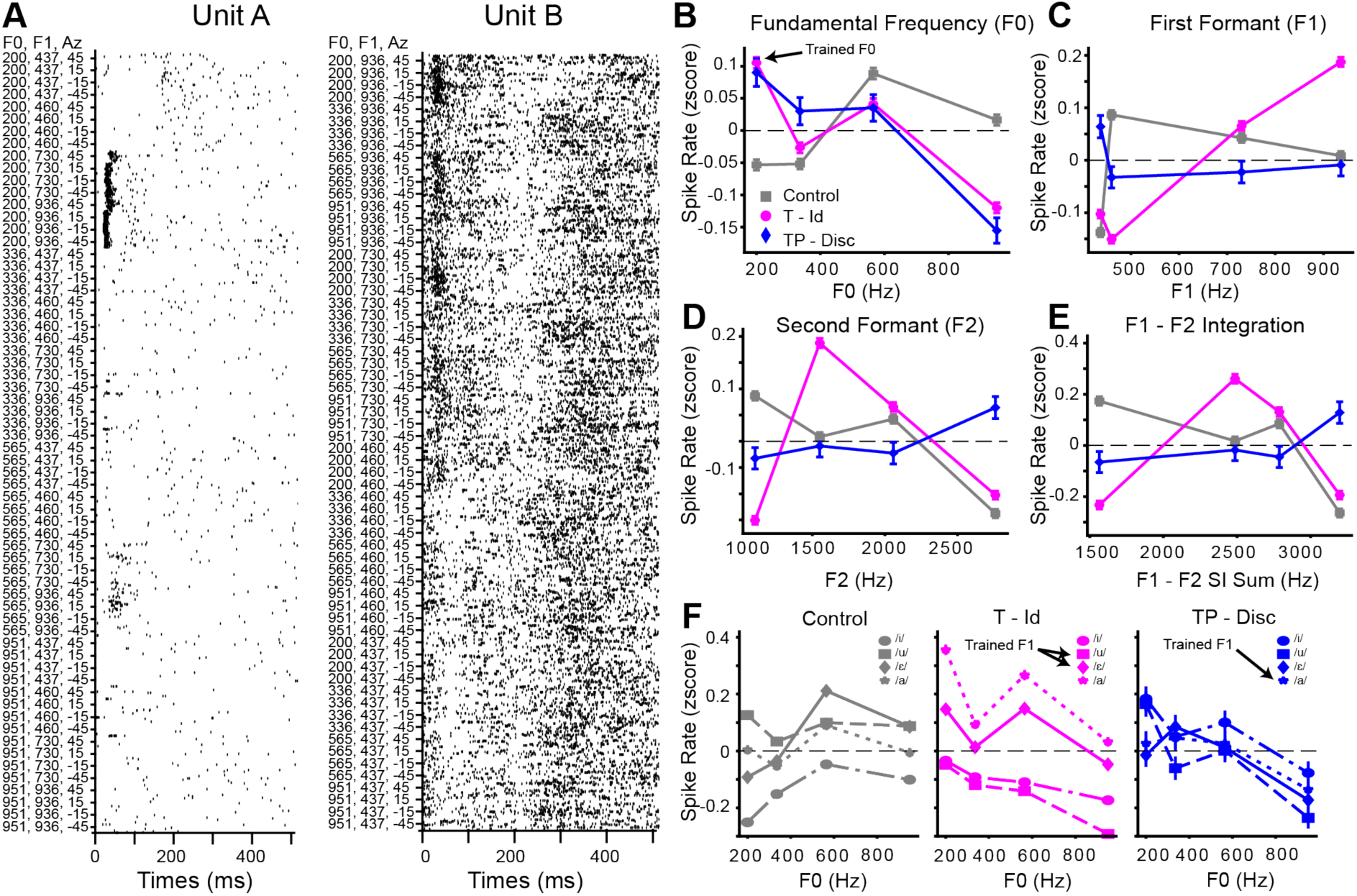
**A** Raster plots from two example units. **B-F** Z-scored spike rates (mean ± SEM,) for all units recorded in the control animals (gray), animals trained in the timbre identification task (T – Id) (magenta), and animals trained in the pitch/timbre discrimination task (TP - Disc) (blue), across F0 (**B**), First formant frequency (F1, **C,** order of vowels from low to high F1:/i/, /u/, /ε/, /a/ ), Second Formant Frequency (F2,**D** order of vowels from low to high F2:/u/, /a/, /ε/, /i/), F1-F2 spectral integration (**E,** order of vowels from low to high: /u/, /a/, /ε/, /i/), F0-F1 (vowels separated and plotted across F0).

While single units appeared to show diverse responses, we first performed some basic analyses to look for any population wide effects of training. To do so, we calculated the normalized (z-scored) spike rate across different stimulus parameters (Figure 2B-F) calculated over the first 200ms after stimulus onset. Both groups of trained animals most frequently heard the stimuli with an F0 of 200 Hz during behavioural testing. We observed larger changes in normalized spike rates for F0 in both training groups than in the control group (Fig. 2B), with both groups of units showing a higher firing rate to sounds with an F0 of 200 Hz than to other F0 values. We examined population level tuning to first formant frequency and second formant frequency (Fig.2C,D) and their integration (Fig.2E, sum of F1 and F2). For timbre the T-ID group in particular showed a much greater modulation of spike rate by timbre, particularly for F1, where higher values to elicited a greater response than lower values. The two trained vowels /u/ (F1 = 460 Hz, F2 = 1105 Hz, trained go left), and / ε / (F1=730 Hz, F2 = 2058 Hz, trained go right) show an enhanced firing rate difference, with neurons appearing to respond more strongly to the contralateral conditioned stimulus and having suppressed responsesfor the ipsilateral trained stimulus. Examination of firing rates across combinations of F0 and timbre (Fig.2F) were suggestive of F0-timbre interactions across all three groups, and in particular the T-id group.

### Response modulation by timbre, pitch and location

Having examined neural tuning at the population level we turned to unit-level analysis using the variance decomposition approaches that we have previously developed (Bizley etl al., 2009) as this allows us to quantify to what extent different perceptual dimensions modulate spiking responses over time. We first asked to what extent neurons were sensitive to each of the three sound features in the trained group as compared to the control group. We determined the proportion of units whose responses were significantly modulated by variation in stimulus location, pitch (determined by fundamental frequency, F0), or timbre. All trained animals learned to discriminate the timbre of the artificial vowels (Fig. 3A, B). Therefore, one might expect to observe a greater number of auditory cortical neurons to convey timbre information post training, as well as fewer neurons that are sensitive to untrained stimulus features (namely, pitch in the T-Id animals and location in both the T-Id and TP-Disc animals).

**Figure 3:**
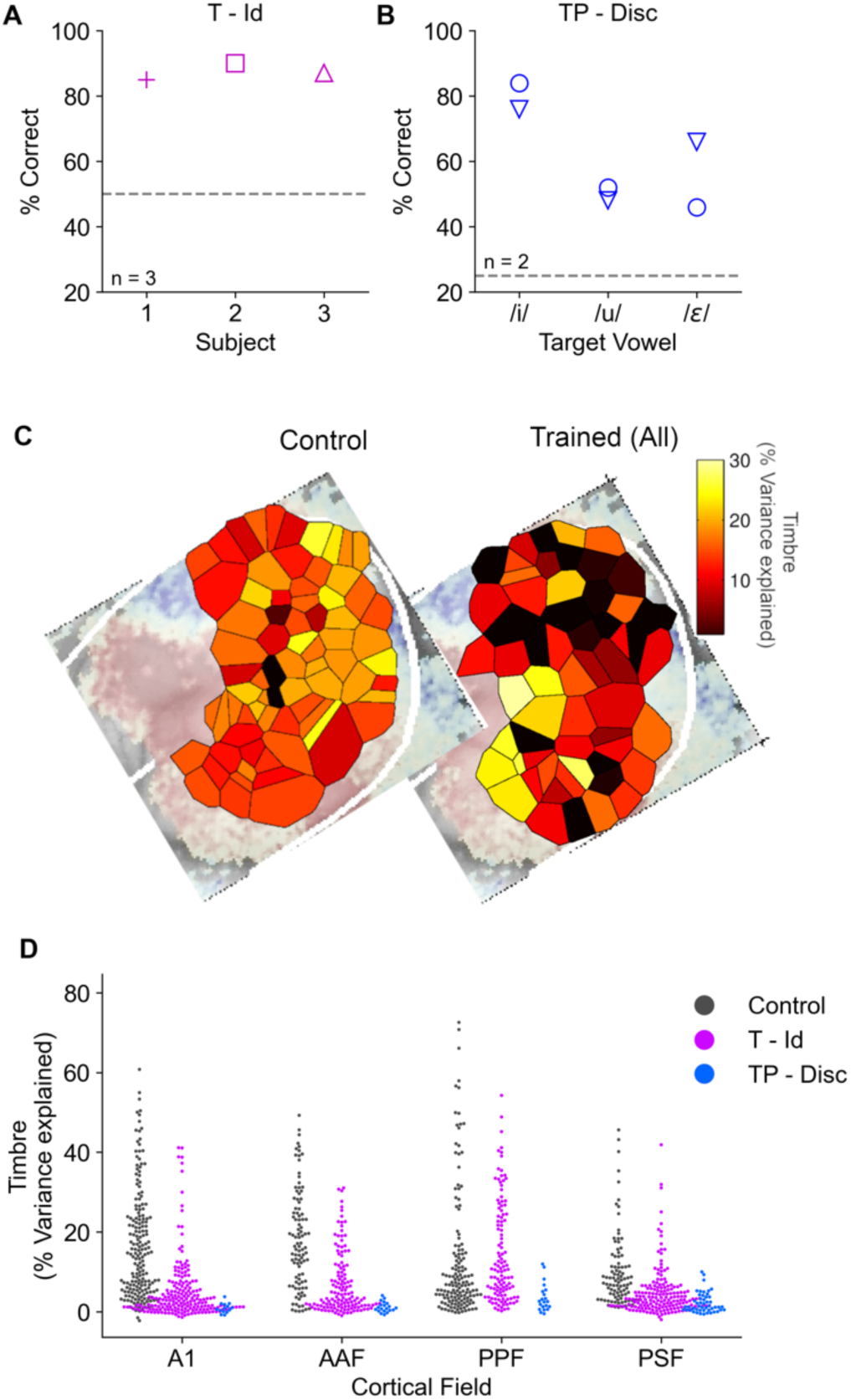
Timbre sensitivity of auditory cortical neurons is reduced in animals trained on timbre discrimination tasks. **A,** Performance of three ferrets identifying timbre across changes in F0 (T-Id, chance = 50%). **B,** Performance of two animals discriminating changes in timbre and F0(TP-Disc, chance = 25%). **C**, Cortical distribution of sensitivity to timbre measured using the proportion of variance explained metric (see Methods). Each tile represents an electrode penetration, with values averaged across all neural units from the same penetration. **D**, Swarm plots showing the distribution of timbre sensitivity across cortical fields and training groups. Each data point is one neural unit.

Contrary to these predictions, the number of units that showed timbre sensitivity was equivalent between trained and control groups (391/783 units and 293/538 units, respectively; *X*^2^ = 0.005, p = 0.95). In both groups, units with joint stimulus sensitivity outnumbered those with sensitivity to only a single parameter, but the distribution of sensitivity to zero, one, two or three stimulus parameters was significantly different between the groups (*X*^2^ = 86.9, p < 0.001). In the trained dataset, many more units showed significant modulation by all three stimulus parameters, or by none of the stimulus parameters, than in the control dataset where most units were modulated by one or two stimulus dimensions (Table 2, example cells in Figure 2A, extended data figure 2.1-5, example rasters number 62, 167, 276, 460, 479. Taken together, this suggests that training resulted in a decrease in the likelihood of units being sensitive to modulations of timbre, pitch or location on their own, but an increase in units sensitive to combinations of all three features. We now consider sensitivity to each feature in turn, asking how the responses of units, and their cortical distribution of sensitivity, varied between trained and control animals.

**Table 2:**
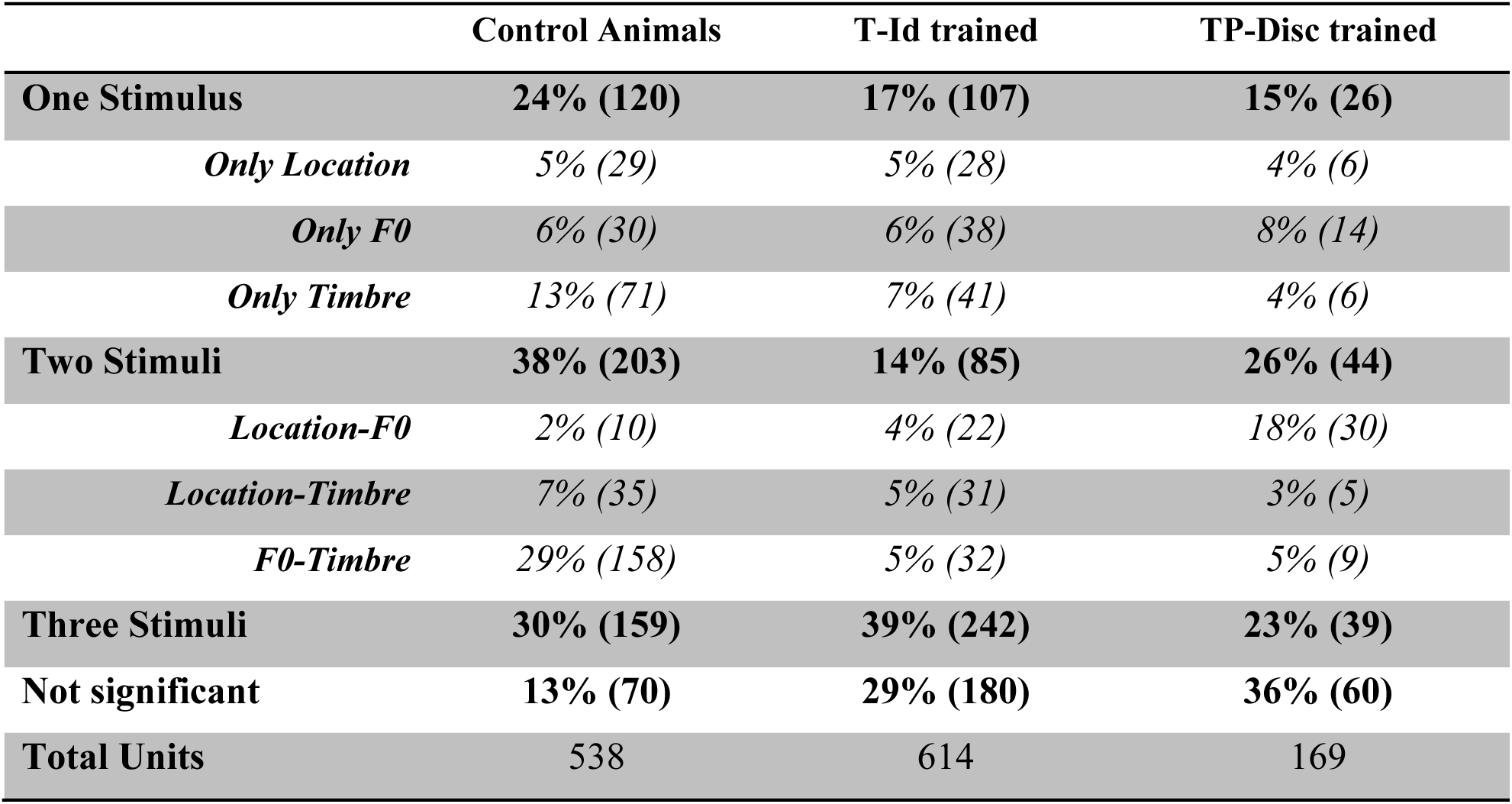
Percentages of units whose responses were significantly modulated by stimulus dimensions timbre, F0, or location, for naïve, T-Id trained and TP-Disc trained animals.

|  | Control Animals | T-Id trained | TP-Disc trained |
| --- | --- | --- | --- |
| **One Stimulus** | **24% (120)** | **17% (107)** | **15% (26)** |
| *Only Location* | 5% (29) | 5% (28) | 4% (6) |
| *Only F0* | 6% (30) | 6% (38) | 8% (14) |
| *Only Timbre* | 13% (71) | 7% (41) | 4% (6) |
| **Two Stimuli** | **38% (203)** | **14% (85)** | **26% (44)** |
| *Location-F0* | 2% (10) | 4% (22) | 18% (30) |
| *Location-Timbre* | 7% (35) | 5% (31) | 3% (5) |
| *F0-Timbre* | 29% (158) | 5% (32) | 5% (9) |
| **Three Stimuli** | **30% (159)** | **39% (242)** | **23% (39)** |
| **Not significant** | **13% (70)** | **29% (180)** | **36% (60)** |
| **Total Units** | **538** | **614** | **169** |

#### Neural sensitivity to trained stimulus features: Timbre

For each unit in our neural dataset, we determined what proportion of the response variance was attributable to each of the three stimulus dimensions, and their combinations. First, we considered sensitivity to timbre, as both behavioral tasks required animals to make judgments based on this feature (Fig. 3A, T-Id; 2B, TP-Disc).

Figure 3C shows the distribution of sensitivity to timbre (i.e. the proportion of variance explained by timbre) mapped across the four cortical fields examined. Although the chi-squared analysis above indicated that the proportion of units with significant timbre sensitivity is equivalent across trained and control groups, the maps in Figure 3C illustrate that significantly less response variance is explained by sound timbre in the trained animals than in the control animals.

We applied a Generalized Mixed Linear Model (GLMM) to assess the impact of different training groups and cortical fields on timbre sensitivity (Table 3) with penetration as a random effect, (in order to account for the shared variability in simultaneous recordings). There was a significant main effect of training, with both the TP-Disc (β = -14.855, p < 0.001) and T-Id (β = -10.816, p < 0.001) training groups showing significant negative effects compared to the control group. The magnitude of these coefficients suggests a marked decrease in the proportion of variance explained by timbre after training. The main effect of cortical field was predominantly decreased timbre sensitivity in field PSF relative to A1 (see Table 3).

**Table 3:**
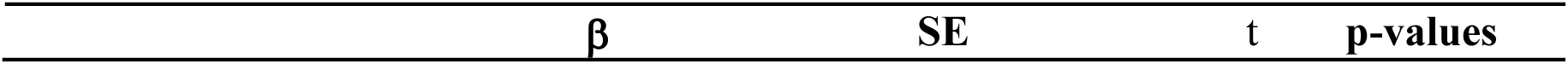
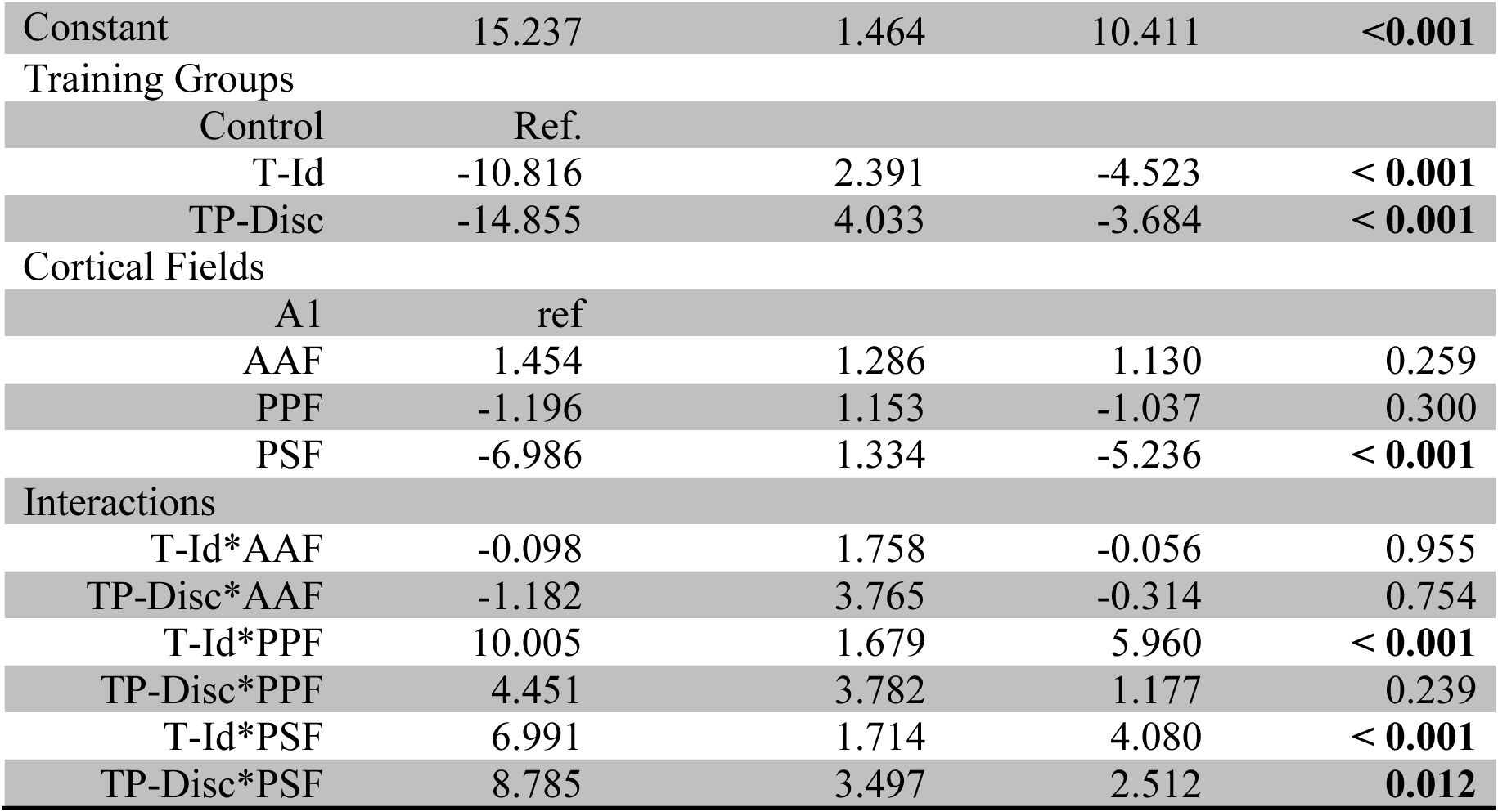
The Generalized Linear Mixed Model results for the proportion of neural response variance explained by timbre. The number of observations was 1265, penetration was included as a random effect. Cross validated root mean squared error was 10.26.

|  |  |  |  |  |
| --- | --- | --- | --- | --- |
| Constant | 15.237 | 1.464 | 10.411 | <0.001 |
| Training Groups |  |  |  |  |
| Control | Ref. |  |  |  |
| T-Id | -10.816 | 2.391 | -4.523 | < 0.001 |
| TP-Disc | -14.855 | 4.033 | -3.684 | < 0.001 |
| Cortical Fields |  |  |  |  |
| A1 | ref |  |  |  |
| AAF | 1.454 | 1.286 | 1.130 | 0.259 |
| PPF | -1.196 | 1.153 | -1.037 | 0.300 |
| PSF | -6.986 | 1.334 | -5.236 | < 0.001 |
| Interactions |  |  |  |  |
| T-Id*AAF | -0.098 | 1.758 | -0.056 | 0.955 |
| TP-Disc*AAF | -1.182 | 3.765 | -0.314 | 0.754 |
| T-Id*PPF | 10.005 | 1.679 | 5.960 | < 0.001 |
| TP-Disc*PPF | 4.451 | 3.782 | 1.177 | 0.239 |
| T-Id*PSF | 6.991 | 1.714 | 4.080 | < 0.001 |
| TP-Disc*PSF | 8.785 | 3.497 | 2.512 | 0.012 |

The GLMM analysis showed several significant interaction terms between group and the secondary fields (Table 3). Therefore, to directly address whether training differentially affected timbre sensitivity in each cortical field, we performed pairwise posthoc comparisons (Tukey’s HSD for multiple comparisons) across groups, separately for each field. These revealed significant decreases in timbre sensitivity in trained animals compared to controls for all cortical fields (p < 0.05) except PPF, which was not significantly different. Overall, these results suggest that long-term training in a spectral timbre discrimination task leads to a field-specific alterations in timbre sensitivity; with the exception of field PPF, trained animals display a decrease in timbre sensitivity throughout tonotopic auditory cortex compared to controls. Example PPF units with high timbre sensitivity can be seen in the extended data Figure 3.1-3: units 327, 550, 677).

While the distribution of CFs was equivalent across fields/groups, we also tested whether the training effects we observed could be explained by differences in the CF distribution, anticipating that low frequency preferring units would also convey more timbre information. We therefore re-ran the GLMM using CF as an additional factor. As expected, CF was negatively associated with timbre sensitivity (high frequency units are less sensitive to timbre than low frequency units) but all of the statistical effects held when timbre was included into the model, including the observation that timbre sensitivity in T-Id animals was preserved in field PPF, but decreased elsewhere (Table 4).

**Table 4:** The Generalized Linear Mixed Model results for the proportion of neural response variance explained by timbre based on training group, cortical field and best frequency.

| Variable | $\beta$ | df | t value | p-value |
| --- | --- | --- | --- | --- |
| **Constant** | 19.085 | 61.288 | 10.713 | <0.001 |
| **Training Group** |  |  |  |  |
| Control | Ref. |  |  |  |
| T-Id | -15.037 | 42.086 | -5.575 | <0.001 |
| TP-Disc | -18.951 | 201.738 | -3.405 | 0.001 |
| **CF** | -0.452 | 960.502 | -3.744 | <0.001 |
| **Fields** |  |  |  |  |
| A1 |  |  |  |  |
| AAF | -6.395 | 1016.637 | -1.837 | 0.067 |
| PPF | -2.521 | 978.265 | -1.338 | 0.181 |
| PSF | -7.708 | 1017.472 | -3.987 | <0.001 |
| **Interactions** |  |  |  |  |
| T – Id*CF | 0.519 | 1020.464 | 2.941 | **0.003** |
| TP – Disc*CF | 0.511 | 1019.011 | 0.908 | 0.364 |
| T – Id*AAF | 4.788 | 1021.544 | 1.196 | 0.232 |
| TP – Disc*AAF | 6.888 | 1003.805 | 0.912 | 0.362 |
| T – Id*PPF | 15.403 | 1018.368 | 6.021 | **<0.001** |
| TP – Disc*PPF | 6.719 | 1018.801 | 1.159 | 0.247 |
| T – Id*PSF | 7.711 | 1013.283 | 3.139 | 0.002 |
| TP – Disc*PSF | 10.090 | 1015.590 | 1.862 | 0.063 |
| CF*AAF | 0.747 | 1016.809 | 3.439 | **<0.001** |
| CF*PPF | -0.027 | 1016.780 | -0.154 | 0.878 |
| CF*PSF | -0.326 | 1021.067 | -1.158 | 0.247 |
| T – Id*CF*AAF | -0.450 | 1016.655 | -1.554 | 0.120 |
| TP – Disc*CF*AAF | -0.866 | 999.553 | -0.543 | 0.587 |
| T – Id*CF*PPF | -0.626 | 1018.538 | -2.245 | **0.025** |
| TP – Disc*CF*PPF | -0.095 | 998.604 | -0.140 | 0.889 |
| T – Id*CF*PSF | 0.277 | 1018.004 | 0.840 | 0.401 |
| TP – Disc*CF*PSF | 0.172 | 1009.431 | 0.255 | 0.799 |

### Formant cues are reweighted in timbre-trained animals

Contrary to our hypothesis, long-term training on a spectral timbre discrimination task led to an overall decrease in neural sensitivity to timbre. However, the stimuli with which we recorded neural responses allowed us to both contrast effects in trained and untrained vowels, and look more generally at tuning to first and second formant frequencies, which our preliminary analysis had highlighted as an area of interest (Fig.2C,D). To contrast responses to trained and untrained vowels, we repeated the variance decomposition analysis using the neural responses to pairs of vowels (i.e. subsets of 50% of the data comprising the responses to stimulus combinations of two vowels presented at each of four F0s and four locations, yielding 32 stimulus conditions in total). The proportion of neural response variance explained by timbre in each of these analyses provided an estimate of how well neuronal responses differentiate a given pair of vowels across variation in F0 and location. We hypothesized that, if training led to enhanced sensitivity to the target vowels, we would observe higher timbre sensitivity (% variance explained) for the trained vowel pairs compared to untrained vowel pairs. Specifically, for the T-Id animals, /u/ versus /ε/ should yield the greatest timbre sensitivity. For the TP-Disc animals, where /a/ was the reference vowel, we expect the /a/-/i/, /a/-/ε/, and /a/-/u/ contrasts to yield higher sensitivity scores than the pairs of vowels that did not include /a/.

We visualized timbre sensitivity across the three groups for all 6 pairs of vowels (Fig.4C), ordering the vowels according to the difference in second formant frequency (ΔF2, Fig.4A-C). The second formant has been shown to strongly influence the perception of trained ferrets when performing vowel discrimination tasks (Town et al., 2015). While the control group showed a clear relationship between timbre sensitivity and ΔF2, such a relationship was not apparent for the trained animals (Fig.4C upper panel). Neither was it the case that timbre sensitivity was enhanced only for trained vowels over untrained ones; or that neurons became exclusively F1 sensitive (Fig.4C lower). To better understand how training group, cortical field, and change in first (ΔF1) and second (ΔF2) formant frequencies influenced the proportion of the neural response variance attributable to timbre, we ran a GLMM that predicted the neural response variance explained by timbre, with factors cortical field, training group, ΔF1 and ΔF2, and with unit ID and penetration as nested random effects. Here, ΔF1 and ΔF2 were calculated as the difference in first and second formant frequencies, respectively, for the relevant vowel pair.

**Figure 4.**
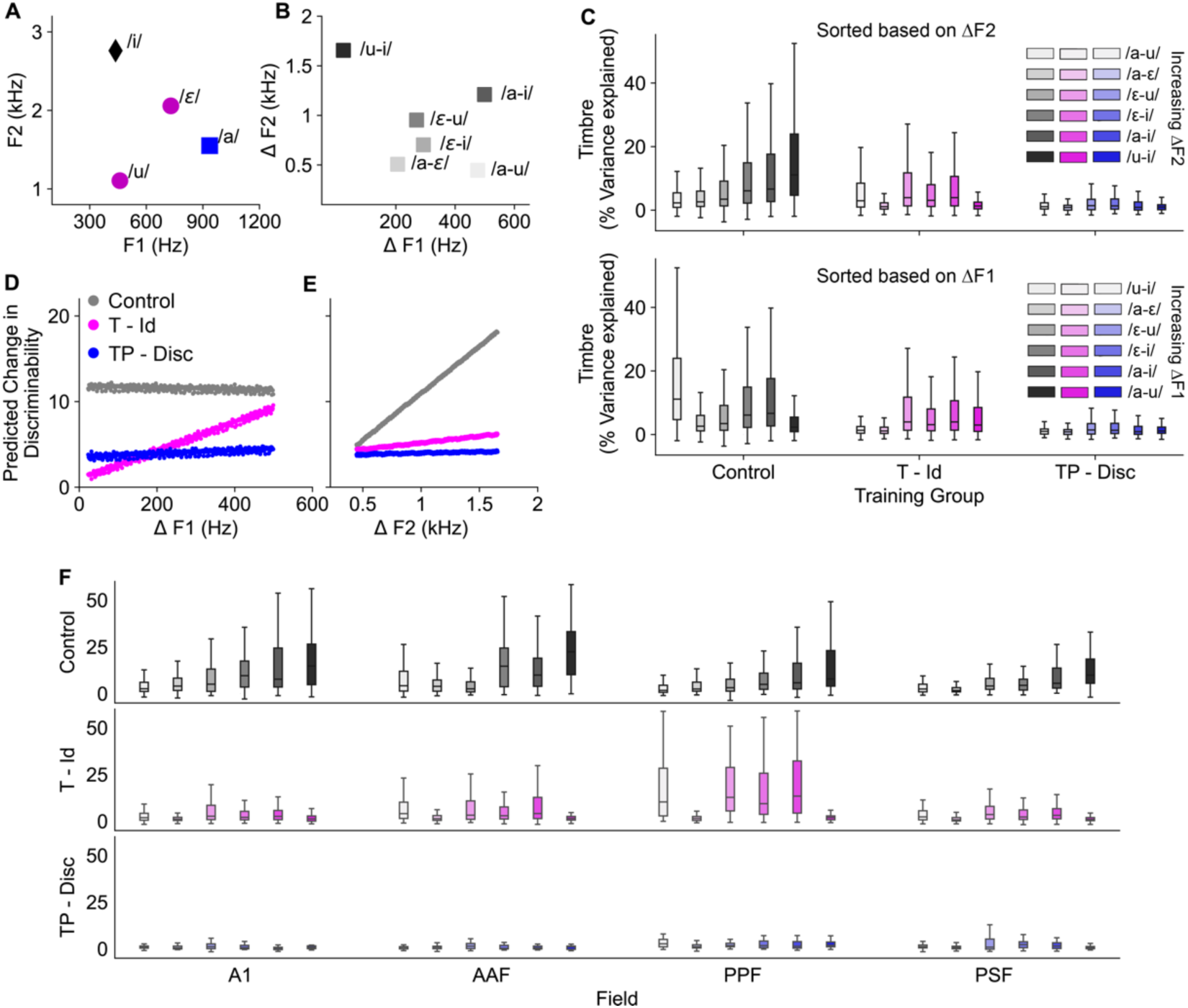
Training re-weights sensitivity to first and second formant frequencies. **A,** First and second formant frequencies for the four vowels used in this study. The /ε/ and /u/ (magenta) was used in T - Id task, and /a/ (blue) was used as reference vowel for TP - Disc task. **B,** Each pair of vowels can be defined according to the difference in their first (ΔF1) and second (ΔF2) frequencies. **C,** The proportion of variance explained by timbre for the six possible vowel combinations, organized according to the magnitude of the difference in second formant frequency (ΔF2, ranked from lowest to highest, top), or first formant frequency (ΔF1 bottom). Error bars are standard error of mean. **D,E,** Model predictions for the change in sensitivity to timbre according to arbitrary differences in F1 (D) and F2 (E). F **F,** The proportion of variance explained by timbre for the six possible vowel combinations (as in C) ordered by ΔF2 and broken down by cortical field.

The results of the GLMM are shown in Table 5, and are consistent with training altering the way in which neurons integrate spectral information. Here positive beta coefficients indicate increased sensitivity (i.e. the model predicts higher amounts of neural response variance explained by timbre). The model showed significant main effects of training group and cortical field (replicating the previous analysis in Table 3 and 4), as well as significant effects of ΔF1 and ΔF2 on sensitivity to timbre, where ΔF1 decreased sensitivity to timbre, and ΔF2 increased sensitivity to timbre. As well as significant main effects, there were multiple significant two-way interactions, all of which were consistent with the trained animals increasing their sensitivity to changes in first formant frequency (e.g. significant ΔF1 x T-Id interaction, with a large positive β coefficient) and and decreasing their reliance on differences in the second formant frequency (significant ΔF2 x T-Id interaction, and ΔF2 x TP-Disc interaction, both with a negative β coefficients). The 3-way interactions were not significant.

**Table 5:** GLMM results for the proportion of variance explained by timbre sensitivity based on training group, cortical field, and differences in first and second formant frequency. The number of observations was 7554, with unit & penetration as nested random effects. Cross validated root mean squared error was 8.65. The three-way interaction encompassing field, formant, and group was not incorporated into the model due to its lack of statistical significance.

| | $\beta$ | SE | t | p-values |
| --- | --- | --- | --- | --- |
| **Constant** | 2.108 | 1.520 | 1.387 | 0.173 |
| **Training Group** |  |  |  |  |
| Control | ref |  |  |  |
| T-Id | 2.361 | 3.013 | 0.783 | 0.440 |
| TP-Disc | 0.518 | 2.681 | 0.193 | 0.849 |
| **Cortical Fields** |  |  |  |  |
| A1 | ref |  |  |  |
| AAF | 0.353 | 1.190 | 0.296 | 0.767 |
| PPF | -3.000 | 1.167 | -2.571 | **0.010** |
| PSF | -4.683 | 1.101 | -4.253 | **< 0.001** |

| Formant Difference |  |  |  |  |
| --- | --- | --- | --- | --- |
| $\Delta F1$ | -0.008 | 0.002 | -3.594 | < **0.001** |
| $\Delta F2$ | 0.010 | 0.001 | 16.026 | < **0.001** |
| Interactions |  |  |  |  |
| TP-Disc*$\Delta F1$ | 0.003 | 0.003 | 1.035 | 0.301 |
| T-Id*$\Delta F1$ | 0.018 | 0.001 | 12.306 | < **0.001** |
| TP-Disc*$\Delta F2$ | -0.011 | 0.001 | -10.539 | < **0.001** |
| T-Id*$\Delta F2$ | -0.010 | 0.001 | -17.390 | < **0.001** |
| AAF*$\Delta F1$ | 0.001 | 0.002 | 0.760 | 0.447 |
| PPF*$\Delta F1$ | 0.001 | 0.002 | 0.691 | 0.490 |
| PSF*$\Delta F1$ | 0.014 | 0.002 | 7.646 | < **0.001** |
| AAF*$\Delta F2$ | <0.001 | 0.001 | 0.673 | 0.501 |
| PPF*$\Delta F2$ | -0.001 | 0.001 | -1.090 | 0.276 |
| PSF*$\Delta F2$ | 0.001 | 0.001 | 1.768 | 0.078 |
| $\Delta F1*\Delta F2$ | < 0.001 | <0.001 | 1.720 | 0.085 |

To further understand timbre sensitivity in trained animals,we used the fitted GLMM coefficients to predict the dependence of timbre-related response variance explained by arbitrary changes in F1 and F2 for both training groups. By systematically varying the ΔF1 and ΔF2 values we simulated a comprehensive sampling of formant space and graphically depicted the resulting predictions for timbre sensitivity (Fig.4D, ΔF1 Fig.4E, ΔF2 visualized). Fig.4G, ΔF1; Fig.4H, ΔF2). The simulated results illustrate the dependence of the control group neurons on ΔF2, and the increased reliance on ΔF1 and decreased sensitivity to ΔF2 present in the T-Id group (see extended data figure 4.1-4 units: 312, 245, 538, 276). In the TP-Disc group, sensitivity to timbre is lower overall, but our limited sample size in these animals prevents clear conclusions from being drawn about the relative dependence on ΔF1 and ΔF2. When dissecting the data by cortical field (Figure 4F), we confirmed that these training-induced shifts in formant frequency weighting were evident across all cortical fields, but are most pronounced in field PPF. We also repeated the GLMM model with CF included as a factor, and the statistical results did not differ (data not shown).

### Sensitivity to fundamental frequency in trained animals

Basic spike rate measures showed that neurons in trained animals fired more when the F0 was 200 Hz (Fig.2B). Since T-Id animals were able to identify trained vowels across a range of randomly varying F0 values (Fig. 5A), we predicted that despite this we might see decreased sensitivity (i.e. increased tolerance) to F0 in the cortical responses of these animals. In contrast, the TP-Disc animals were not required to generalize their discriminations across a second stimulus feature, but were instead trained to detect changes in either F0 (Fig. 5B) or spectral timbre (Fig. 3B). Therefore, we expected to potentially see greater F0 sensitivity in the cortical responses of these animals, compared to the T-Id and control groups. While some units were clearly tuned to F0 (e.g. Extended data Figure 5.1-2, unit: 92, and 276, both tuned to F0=336Hz) and others were driven most strongly by 200Hz F0 stimuli (e.g. Extended data Figure 5.3-5, units: 362, 309, 363), when sensitivity to F0 is plotted across the cortical surface (Fig. 5C), or broken down by cortical field (Fig. 5D), it is apparent that both trained groups show decreased F0 sensitivity compared to the control group.

**Figure 5:**
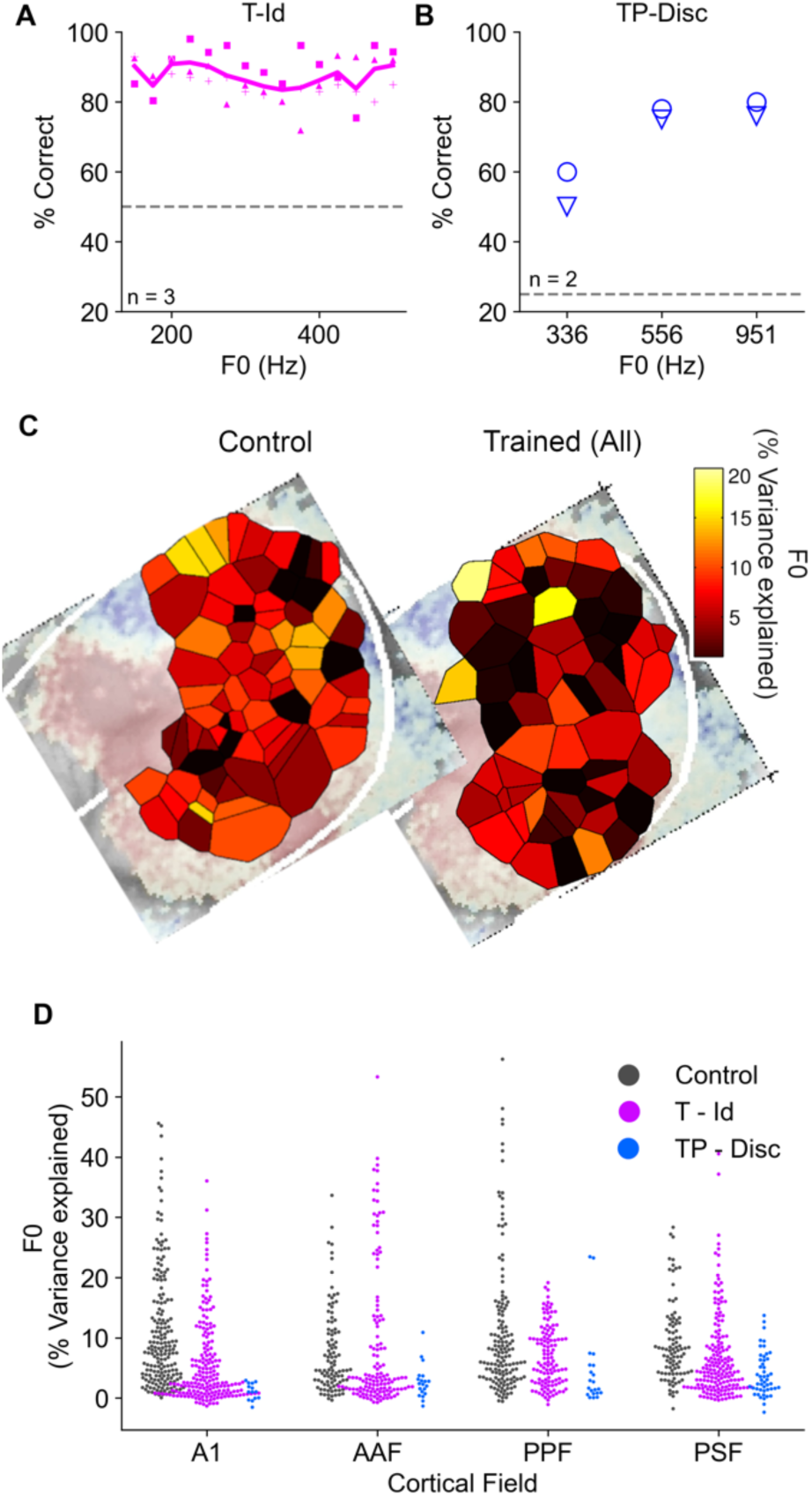
Cortical sensitivity to F0 is decreased in trained animals. **A,** Behavioral performance for discriminating timbre across fundamental frequency (F0) on the timbre identification task (T-Id). **B,** F0 timbre discrimination performance for two animals (TP-Disc). Symbols show individual animals. **C,** Voronoi tessellation maps plotting the proportion of variance explained by F0 for control (left) and trained (right) animals. Conventions as in Figure 3C. **D,** Swarm plots showing the distribution of F0 sensitivity across neuron units in different cortical fields and training groups.

We employed a GLMM to assess the influence of auditory training and cortical field on neural sensitivity to F0 (Table 5). The results indicated that both TP-Id and T-Disc groups exhibited a significant reduction in F0 sensitivity compared to the control group, with respective coefficients of β = -11.246 and β = -6.330 (both p < 0.001). In terms of cortical fields, a marked decrease in sensitivity was observed in AAF (β = -3.585, p < 0.001) relative to A1, while PSF and PPF were not significantly different from A1. There were significant interaction effects between training groups and cortical fields. These findings run counter to our hypothesis that F0 sensitivity in auditory cortex would be enhanced by F0-specific training (i.e., the TP-Disc group) and decreased in animals trained to ignore F0 changes (i.e., the T-Id group). Instead, we observed an overall reduction in neural sensitivity to F0 in trained animals.

**Table 5.** The GLMM results for the proportion of variance explained by F0 sensitivity. The number of observations was 1265, with penetration as nested random effects. Cross validated root mean squared error was 8.33.

| | $\beta$ | SE | t | p-values |
| --- | --- | --- | --- | --- |
| Constant | 11.160 | 0.799 | 13.964 | < 0.001 |
| Training Groups |  |  |  |  |
| Control | ref |  |  |  |
| T-Id | -6.330 | 1.254 | -5.050 | < 0.001 |
| TP-Disc | -11.246 | 2.610 | -4.309 | < 0.001 |
| Cortical Fields |  |  |  |  |
| A1 | ref |  |  |  |
| AAF | -3.585 | 1.024 | -3.501 | < 0.001 |
| PPF | -0.566 | 0.915 | -0.618 | 0.536 |
| PSF | -2.030 | 1.057 | -1.919 | 0.055 |
| Interactions |  |  |  |  |
| T-Id*AAF | 7.065 | 1.391 | 5.081 | < 0.001 |
| TP-Disc*AAF | 5.188 | 3.017 | 1.719 | 0.086 |
| T-Id*PPF | 1.090 | 1.337 | 0.815 | 0.415 |
| TP-Disc*PPF | 5.125 | 3.004 | 1.706 | 0.088 |
| T-Id*PSF | 3.098 | 1.363 | 2.273 | 0.023 |
| TP-Disc*PSF | 5.963 | 2.757 | 2.163 | 0.031 |

### Sensitivity to an untrained feature: Location

Finally, we examined how training impacted cortical sensitivity to sound source location cues. In all cases all behaviorally relevant sounds were presented from a speaker in front of the central nose-poke sensor. In our control dataset, we previously observed that the spatial tuning of neurons in response to these vowel stimuli presented in virtual acoustic space was modest, and as expected, predominantly contralateral (Bizley et al., 2009). However, when the same neurons were tested with spatially modulated broadband noise (also in virtual acoustic space), we observed considerably greater spatial modulation, leading us to conclude that the low spatial sensitivity was a likely product of the limited bandwidth of the artificial vowel stimuli. Here, we examined whether exposure to the vowels in the behavioral task altered the spatial sensitivity measured in auditory cortex using these same stimuli. As Figure 6 shows, spatial sensitivity was indeed higher in trained animals, which can be visualized across the auditory cortical surface (Fig.6A) or between cortical field (Fig.6B). Specifically, in the non-primary cortical areas, spatial sensitivity appeared to be higher in the T-Id animals than in controls.

**Figure 6:**
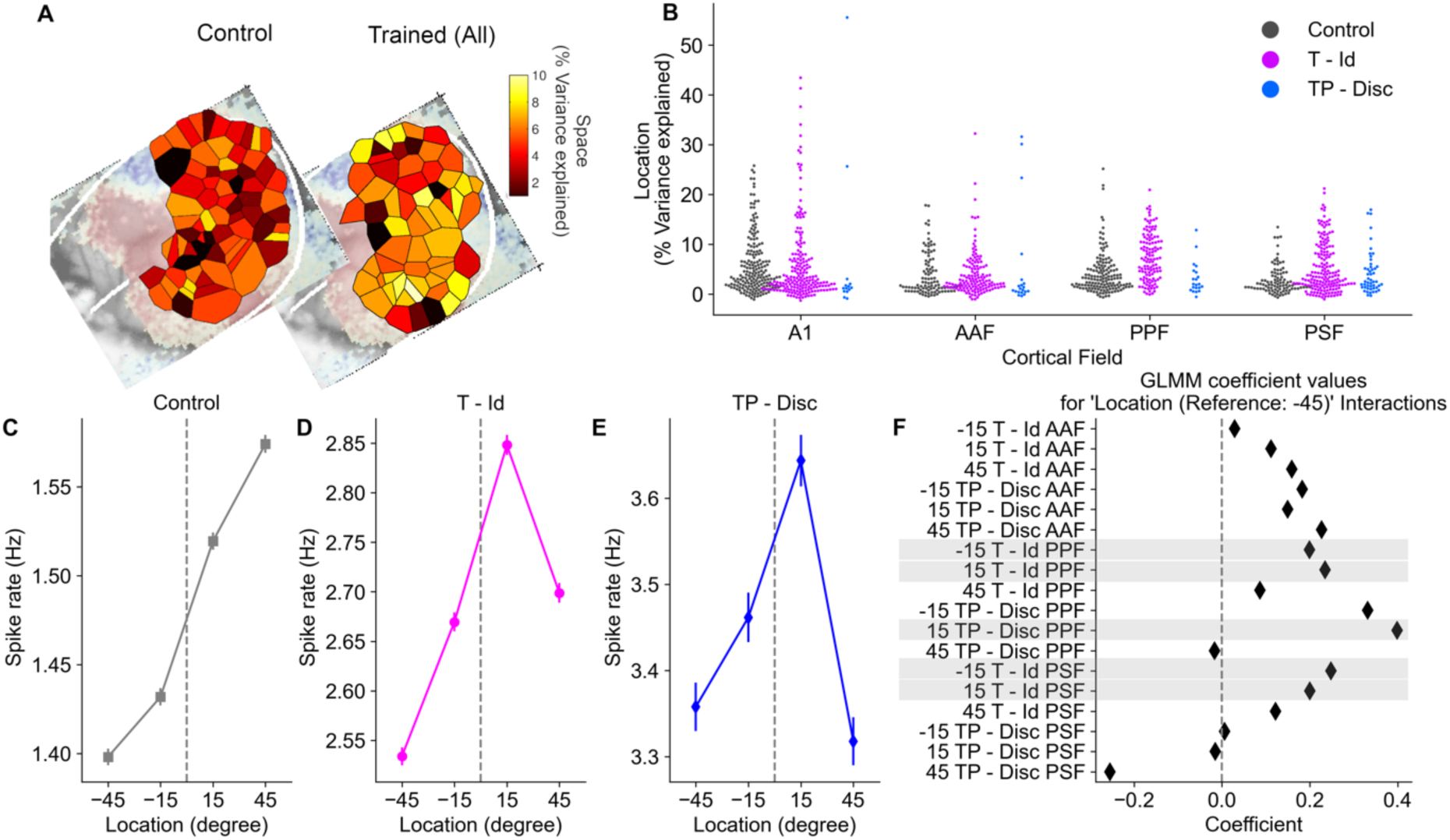
Cortical sensitivity to location, an untrained feature, is higher in trained animals. **A**, Voronoi tessellation maps plotting the proportion of variance explained by location for control and trained animals. **B**, Swarm plots showing the proportion of variance explained by location across fields and training groups. **C-E**, Population spatial tuning functions (mean ± SEM for all responsive units) for control, T-Id animals and TP-Disc animals. **F,** Beta coefficients for the impact of GLMM parameters on spike rate, parameters showing significant effects (p < 0.05) are emphasized with gray bars.

To examine whether training altered the neural sensitivity to sound source location we ran a GLMM using training group, cortical field and their interactions as factors (Table 7). There was no significant overall effect of group on spatial sensitivity, but there were significant effects of cortical field, with all areas showing lowered spatial sensitivity relative to A1 (See Table 7 for details). There were also significant interactions (with positive β coefficients) for training group and fields PSF and PPF for the T-Id animals, supporting the observation that azimuth sensitivity was higher in the non-primary fields in trained animals.

**Table 7:** The GLMM results for the proportion of variance explained by sound source location sensitivity. The number of observations was 1265, with penetration as nested random effects. Cross validated root mean squared error was 5.53.

| | $\beta$ | SE | t | p-values |
| --- | --- | --- | --- | --- |
| Constant | 5.664 | 0.586 | 9.659 | < 0.001 |
| Training Groups |  |  |  |  |
| Control | ref |  |  |  |
| T-Id | -0.083 | 0.931 | -0.089 | 0.930 |
| TP-Disc | -0.237 | 1.856 | -0.128 | 0.899 |
| Cortical Fields |  |  |  |  |
| A1 | ref |  |  |  |
| AAF | -2.178 | 0.682 | -3.196 | **0.001** |
| PPF | -1.676 | 0.603 | -2.777 | **0.006** |
| PSF | -3.183 | 0.708 | -4.497 | **< 0.001** |
| Interactions |  |  |  |  |
| T-Id*AAF | -0.103 | 0.925 | -0.111 | 0.912 |
| TP-Disc*AAF | 0.951 | 1.989 | 0.478 | 0.633 |
| T-Id*PPF | 2.544 | 0.888 | 2.866 | **0.004** |
| TP-Disc*PPF | -0.503 | 1.985 | -0.253 | 0.800 |
| T-Id*PSF | 2.426 | 0.906 | 2.679 | **0.008** |
| TP-Disc*PSF | 1.879 | 1.863 | 1.009 | 0.313 |

We next asked whether the increased spatial sensitivity in trained animals reflected altered spatial tuning. To assess this, we constructed spatial response functions for each neuron, taken as the mean sound-evoked spike rate at each location across all pitch and timbre combinations. To derive a population spatial receptive field, we averaged these functions across all recorded units (Fig.6C-E). As expected, in the control animals (Fig.6C), this yielded a monotonically increasing function with the most contralateral (+45°) location eliciting the strongest firing rates. In contrast, in both trained groups of animals, spatial response profiles were non-monotonic and showed a peak at +15° azimuth. This midline tuning was also clearly evident in the raw data (Extended data figure 6.1-3, units: 185, 82, 309).

The GLMM in Table 7 showed significant group x field interactions, suggesting that training effects on spatial sensitivity differ across cortical fields. To quantify these effects, we ran a GLMM to model spike rates with group (T-Id or TP-Disc relative to the reference category control group), cortical field (reference category A1), and spatial position (categorical predictor, relative to -45°) as factors, with penetration and unit as random effects. This confirmed significant main effects of spatial position (i.e. responses at -45° and +45° azimuth were significantly different, reflecting the dominant contralateral tuning of the units), and significant field*group*position interactions for PPF and PSF in both trained groups (Fig.6F). The coefficients for these effects were negative for +45° and positive for ±15°, confirming the changes observed in the azimuth-response plots (Fig.6C-E). Thus, although sound source location was not varied during behavioral testing, and was not relevant to the task, the representation of the region from which the sounds were presented (i.e. the midline) was enhanced in the trained animals.

## Discussion

In this study, we trained two groups of ferrets to discriminate perceptual attributes of artificial vowels. One group (T-Id animals) categorized vowels according to their identity (i.e. spectral timbre) across changes in pitch (i.e. F0), while the other was trained to detect changes in either the F0 or timbre in a sequence of on-going vowel sounds (TP-Disc animals). We predicted that the T-Id animals would show enhanced neural sensitivity for timbre and increased tolerance (i.e. decreased sensitivity) to other sound features. In fact, what we observed was more complex: sensitivity to timbre was lower in three auditory cortical areas of trained animals compared to controls, but was maintained in PPF (a secondary tonotopic region). Moreover, cortical sensitivity to vowel identity was less contingent on changes in the frequency of the second formant in trained animals, and instead was instead dependent on changes in both the first and second formant frequency. Sensitivity to F0, which the T-Id animals were trained to generalize across, decreased over all fields with some evidence for a field specific increase in PSF. In contrast, sensitivity to sound source location, which did not vary during the tasks, was enhanced in non-primary fields of trained animals. Furthermore, spatial receptive fields were shifted towards the midline, from where the target sounds originated. Overall, animals trained to discriminate vowels in both tasks showed an unexpected decrease in cortical sensitivity to timbre and F0 relative to controls, and enhanced spatial sensitivity.

Previous studies investigating the impact of training on neural tuning in auditory cortex have focused primarily on map plasticity in A1(Irvine, 2018). However, neurons in higher auditory cortical fields are thought to become increasingly specialized for processing spatial or non-spatial stimulus attributes (Rauschecker and Tian, 2000; Bizley and Cohen, 2013; Elgueda et al., 2019), and show enhanced attention-related changes in activity during behavior (Mesgarani and Chang, 2012; Atiani et al., 2014; Elgueda et al., 2019). While most studies of the cortical substrates for perceptual learning have focused on A1, our data suggest that the areas that show larger attention-related changes may also be those in which receptive fields are optimized through learning to process task-relevant stimuli.

The observation of a marked decrease in sensitivity to task-relevant features in the primary auditory cortical fields A1 and AAF is a potentially surprising finding. However, a number of studies report that engagement in a behavioral task causes suppression of neural responses in auditory cortex (Otazu et al., 2009; Town et al., 2018) and training to discriminate natural sounds has been observed to result in sparser auditory cortical representations of these sounds in A1 of mice (Maor et al., 2019). The reduced sensitivity we see, along with the increase in unresponsive neurons, may reflect a sparsening of representations. Additionally, it may result from the integration of diverse non-sensory inputs into auditory cortex that ultimately underlie the emergence of choice- or motor-related activity. Cooling A1 in ferrets does not lead to an impairment in a vowel discrimination in a task analogous to the one used here, with behavioural deficits only evident when vowels are presented in simultaneous noise (Town et al., 2023). Our neural data raise the testable prediction that inactivation of PPF may cause a timbre identification deficit, whereas cooling A1, AAF or PSF would have a more modest effect on performance in these tasks.

Training ferrets oto identify vowels altered the sensitivity of cortical neurons to the cues that underlie spectral timbre. While the complex sounds used here may allow animals to use a variety of acoustical criteria to solve the tasks, individual animals use remarkably consistent strategies to identify vowel timbre, even across different cohorts, laboratories, and stimulus conditions (Bizley et al., 2013; Town et al., 2015). In the naïve control animals, the most discriminable stimuli were those in which there was a large difference in F2. This finding is mirrored behaviorally; when first and second formant cues are placed in conflict, ferrets and human listeners tend to weight the position of the second formant over the first in their decisions, with their behavior being best predicted by either F2 position or the position of the spectral centroid (Town et al., 2015). Nonetheless, the T-Id animals described here were tested behaviorally with single-formant stimuli and they were able to accurately classify F1 for /u/ and F2 for /ε/ (Bizley et al., 2013). <u>The finding that neural responses in the trained animals are</u> <u>explained by changes in both the first and second formants for trained and novel vowels mirrors</u> <u>electrophysiological recordings in humans (Oganian et al., 2023) and suggests that learning</u> <u>may be associated with enhanced integration of the cues that define the spectral envelope, and</u> <u>an increased sensitivity to low frequency spectral peaks.</u> While the highest levels of sensitivity to timbre were observed in PPF of trained animals, the pattern of cue-weighting was preserved across fields, suggesting that it is widespread within the auditory cortex. This finding is consistent with previous studies in A1 (Keeling et al., 2008; Beitel et al., 2020). At the population level, control animals showed a reasonably flat firing rate distribution across vowels. In contrast the T-ID animals showed much greater modulation with a reduced firing rate for one trained vowel (/u/, conditioned as “go left”) than the other (/ρ,/, conditioned as “go right”). Given our recordings were in left auditory cortex, this pattern is consistent with recent reports that in animals trained in 2AFC experiments tuning within a hemisphere of A1 is optimized for the contralateral stimulus (Znamenskiy and Zador, 2013; Chang et al., 2022), but confirmation of this would require bilateral recordings.

Artificial vowels are low frequency stimuli (<4 kHz). Our sampling yielded balanced samples of neurons across the frequency axis in both trained and control animals (Fig.1). Although we did not design our experiments to perform high-resolution within-field mapping, our data show no indication of an increase in the proportion of low-frequency recording sites suggestive of tonotopic reorganization after training. We also considered the impact of training on the neural representation of two other sound features: In both groups of trained animals there was a decrease in F0 sensitivity, despite the fact that animals in the T-Id task were required to generalize timbre decisions across F0, while the TP-Disc animals actively discriminated both pitch and timbre features. Our sampling approach ensured that we had good coverage in each cortical field, thereby facilitating comparisons between them. By opting to sample multiple cortical areas, however, it is possible that we did not have sufficient resolution to detect possible hot-spots of pitch sensitivity, such as the “pitch area” proposed by Bendor and Wang based on data from the marmoset (Bendor and Wang, 2005), but which has not yet been corroborated in other mammalian species.

In contrast to the decreased sensitivity to non-spatial sound features, changes in spatial sensitivity in trained animals despite the task having no spatial component to it. These changes occurred principally as an increase in spatial sensitivity in the non-primary fields PPF and PSF and resulted in population tuning shifts from monotonically increasing for more contralateral sounds to peaking 15° contralateral to the midline. Our spatial receptive fields were coarsely measured with stimuli at ±15° and ±45°, but if the change in tuning observed in the recorded hemisphere were mirrored on the other side of the brain, this should result in an expanded representation of the midline, where stimuli were presented during behavioral testing. Engaging in a sound discrimination task has been shown to refine spatial tuning in cat A1, with changes occurring for both spatial and – more modestly – non-spatial tasks (Lee and Middlebrooks, 2011). While previous studies suggest active discrimination of the trained feature is not necessary to drive changes in neural responses (Keeling et al., 2008) this is, to our knowledge, the first report of enhanced location coding after repeated exposure to behaviorally-relevant stimuli. Furthermore, these changes were observed primarily in higher- order auditory cortex (PPF and PSF), rather than A1.

A limitation of our study is the small sample size in the TP-Disc group. However, despite the limited statistical power, and differences in the behavioural task design, the data from these animals support the main conclusions from the T-Id group; namely diminished timbre sensitivity overall, with an increased sensitivity to combinations of features and down- weighting of sensitivity to delta-F2, and a shift in spatial tuning to midline locations. A further caveat is that the recordings were performed under anesthesia, which may underestimate neuronal changes(Chang et al., 2022). This was essential for performing mapping unit activity across multiple neighboring cortical fields in the same animals.

Methods for recording in awake animals currently allow only high-density sampling from a small area of cortex, or sparse sampling across multiple fields. Recording under anesthesia also has the advantage of allowing us to separate out effects of sensitivity to stimulus features from attention and to measure the receptive field features to which attentional and task- related effects are likely added during active listening. Tuning for higher-order perceptual features such as pitch and timbre, may also be modulated by attention during task engagement, so it would be interesting to examine this form of short-term plasticity in future studies, to help us better interpret the relevance of the long-term plasticity investigated here.

In summary, training ferrets to discriminate or detect specific features present in artificial vowels has diverse effects on the sound stimulus sensitivity of neurons in auditory cortex. Sensitivity to task-relevant stimulus features, which is broadly distributed in naïve animals, is modulated by training in a manner that differs across auditory cortical fields. Furthermore, in contrast to control animals, in which responses of cortical neurons are strongly weighted by the second formant frequency of artificial vowels, the units recorded in trained animals integrate information about both the first and second formant frequency. Finally, sensitivity to a task-orthogonal feature – here auditory space – is enhanced when training stimuli are repeatedly presented from a single location. Since learning triggers widespread changes in gene expression in auditory cortex (Graham et al., 2023), future work can seek to unpick the specific molecular mechanisms that support changes in auditory cortical function and auditory memory, and consider regions beyond auditory cortex that might support or drive such changes (Jia et al., 2024).

## Acknowledgments

This work was supported by a RNID (formally Action on Hearing Loss) PhD studentship to H.A., grants from the Biotechnology and Biological Sciences Research Council (grants BB/D009758/1 to J.W.H.S., J.K.B., and A.J.K. and BB/H016813/1 to J.K.B.), a Royal Society Dorothy Hodgkin Fellowship (J.K.B.), a Sir Henry Dale Fellowship (J.K.B., WT098418MA) and a Wellcome Principal Research Fellowship (A.J.K., WT108369/Z/2015/Z).

