## Extended data for "Auditory training alters the cortical representation of complex sounds"

Extended data
Figure 2-1

cellID: 92, Animal: 833, Field: PSF
TP - Disc Trained

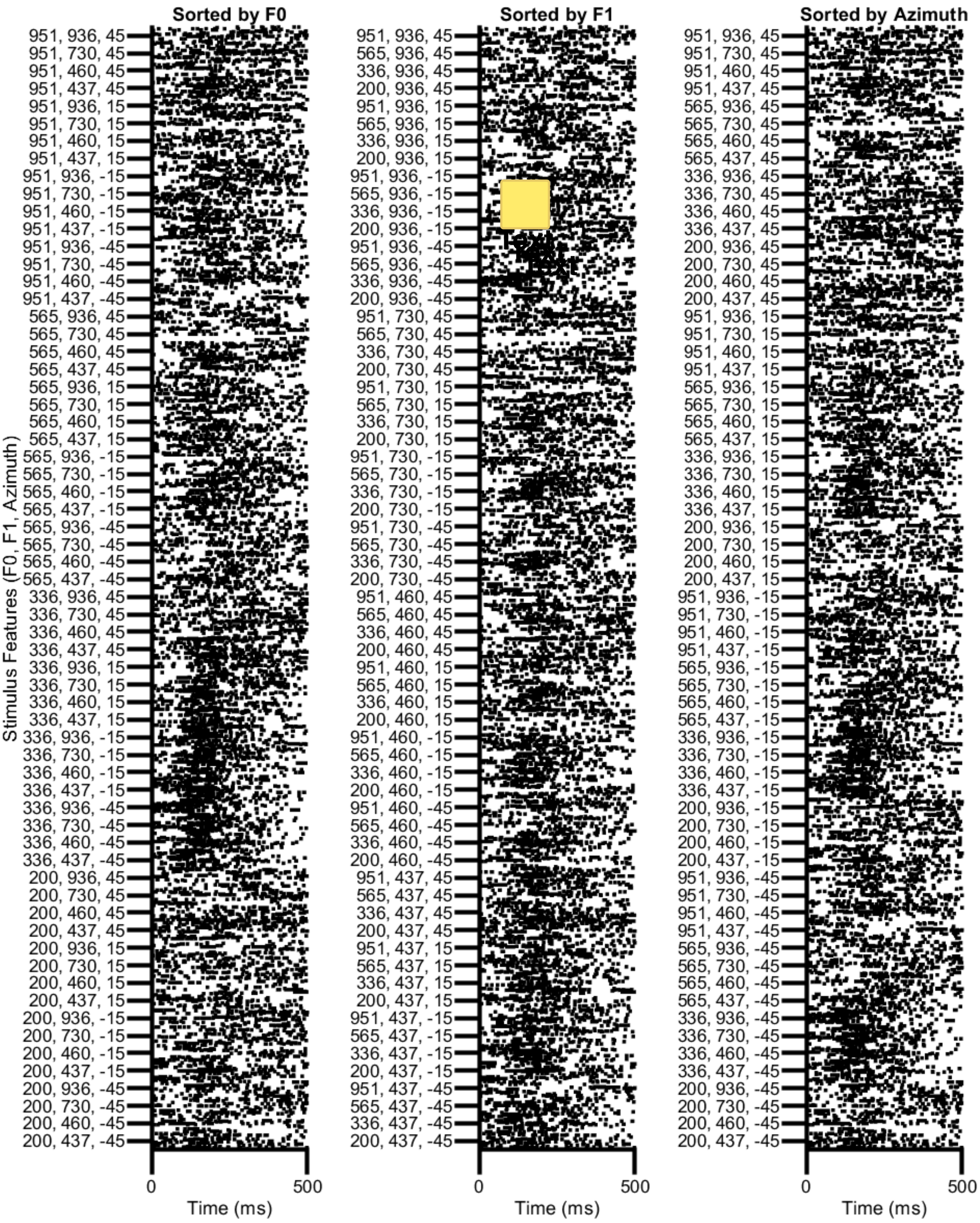

Extended data 2-2 Extended data 4-1

cellID: 167, Animal: 848, Field: AAF
TP - Disc Trained

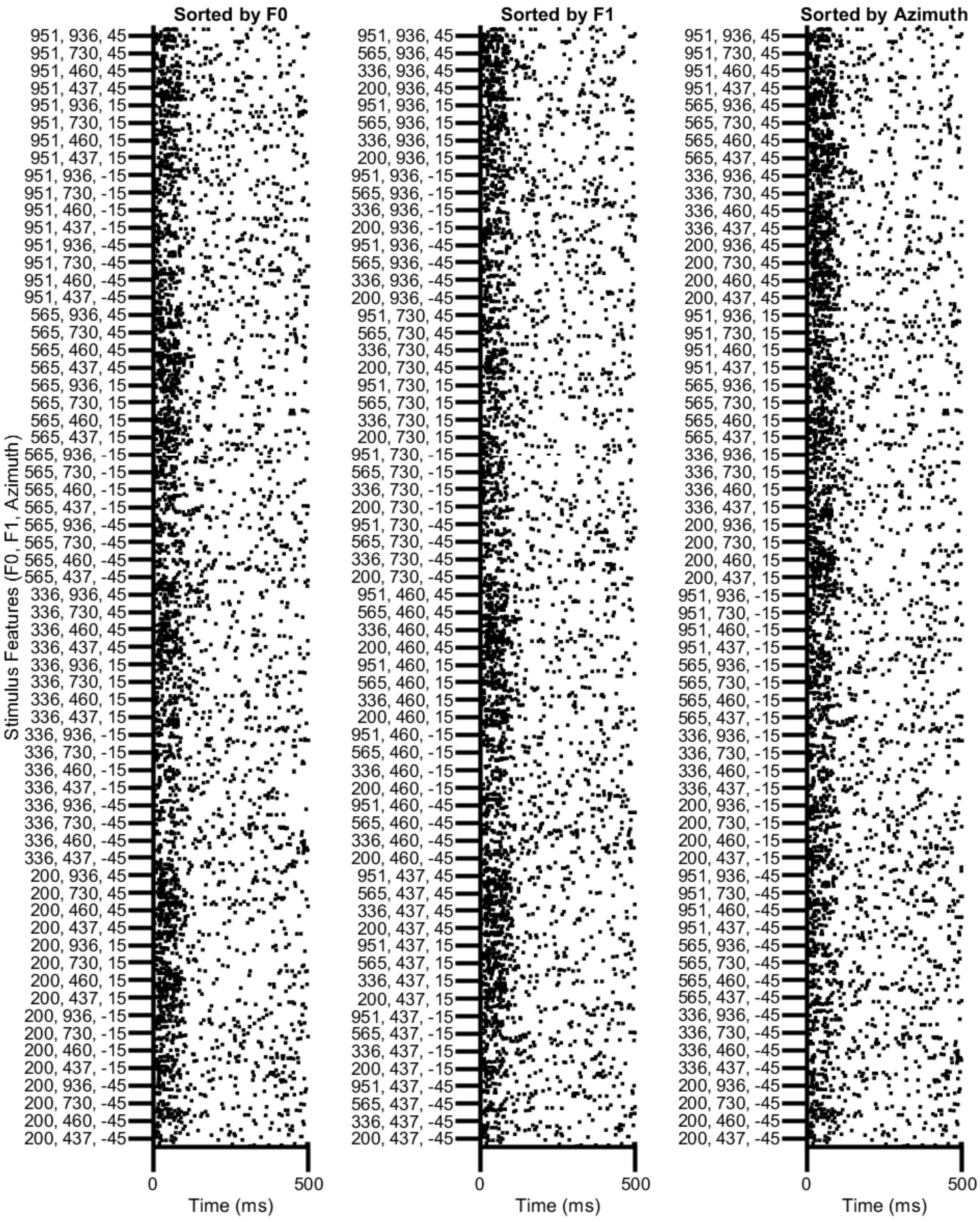


xtended data 2-2

cellID: 276, Animal: 943, Field: PSF
T - ld Trained

Type text here

Extended Exdata 2-3

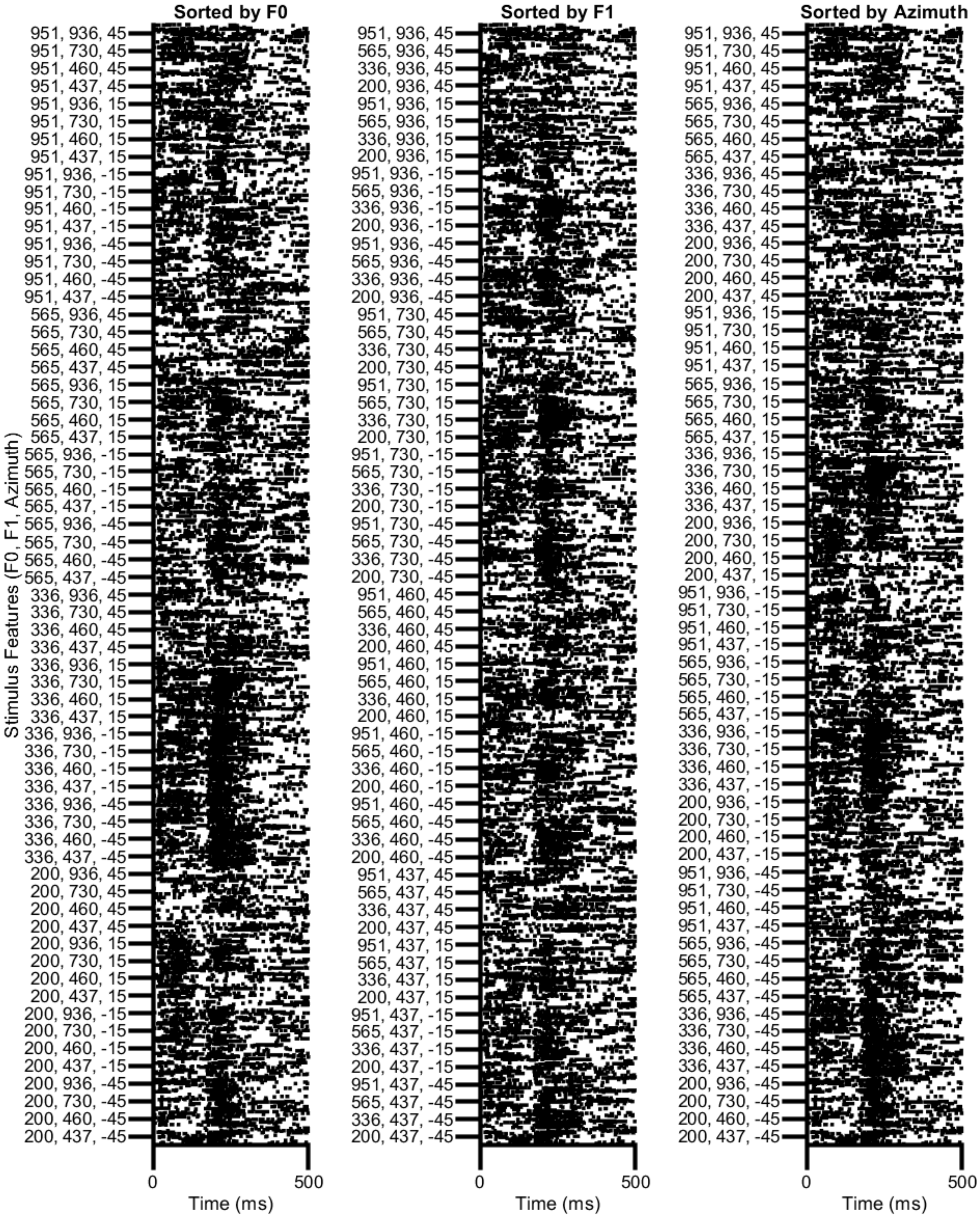

Extended data 2-4

cellID: 460, Animal: 943, Field: AAF
T - ld Trained

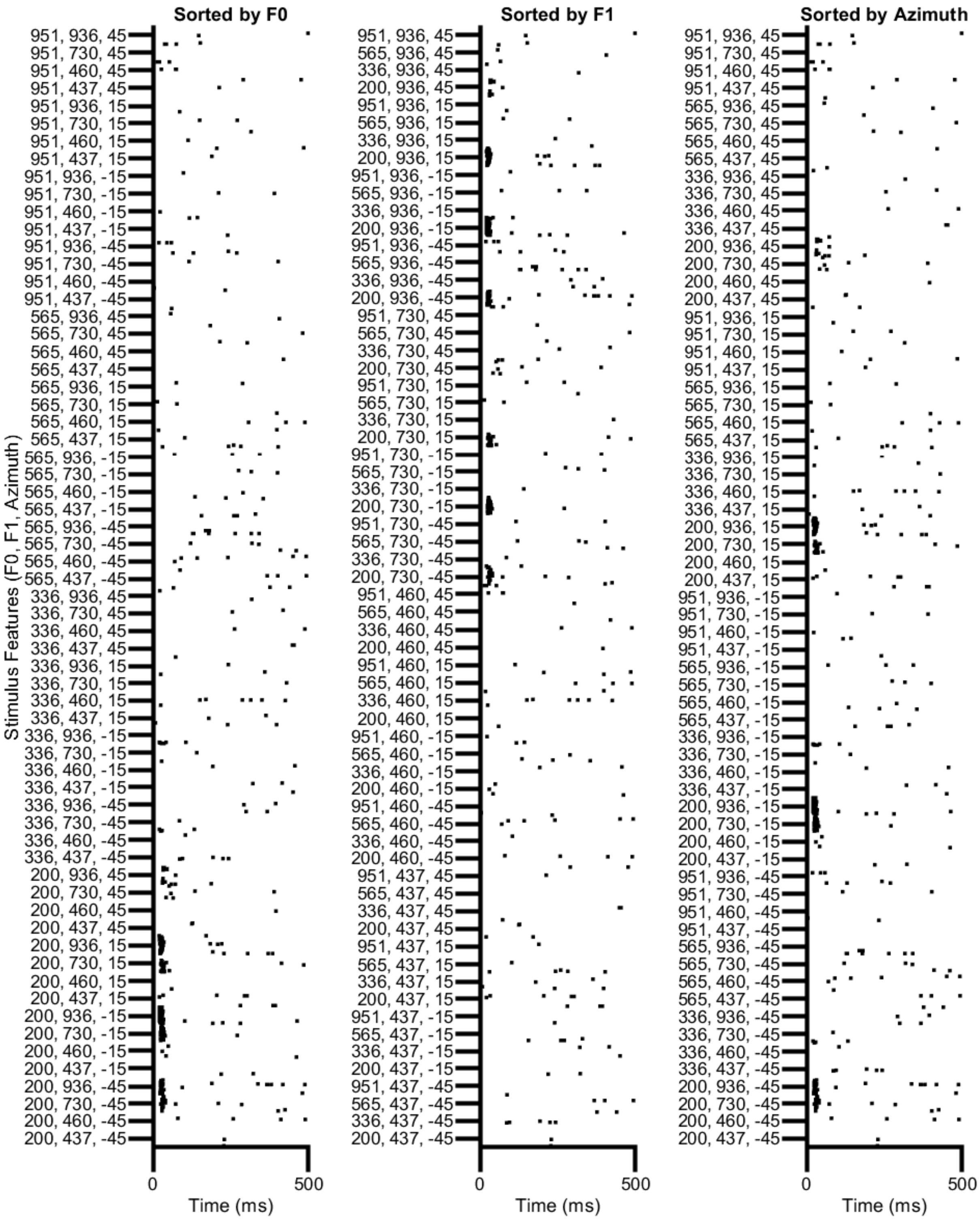

Extended data 2-5

cellID: 479, Animal: 963, Field: AAF
T - ld Trained

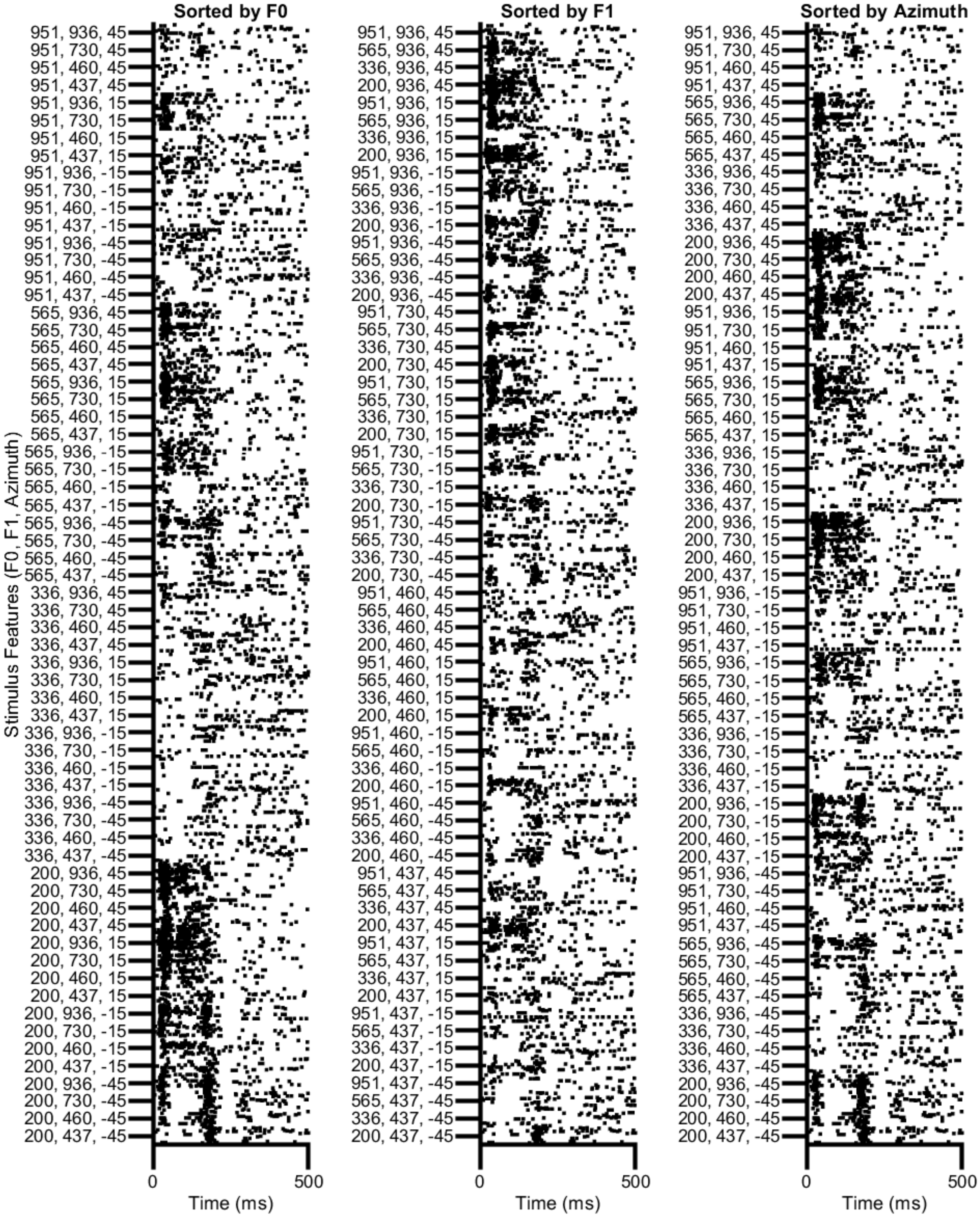


=ended data 2-2

Extended data 3-1

### cellID: 327, Animal: 943, Field: PPF
T - Id Trained

Extended data 6-3

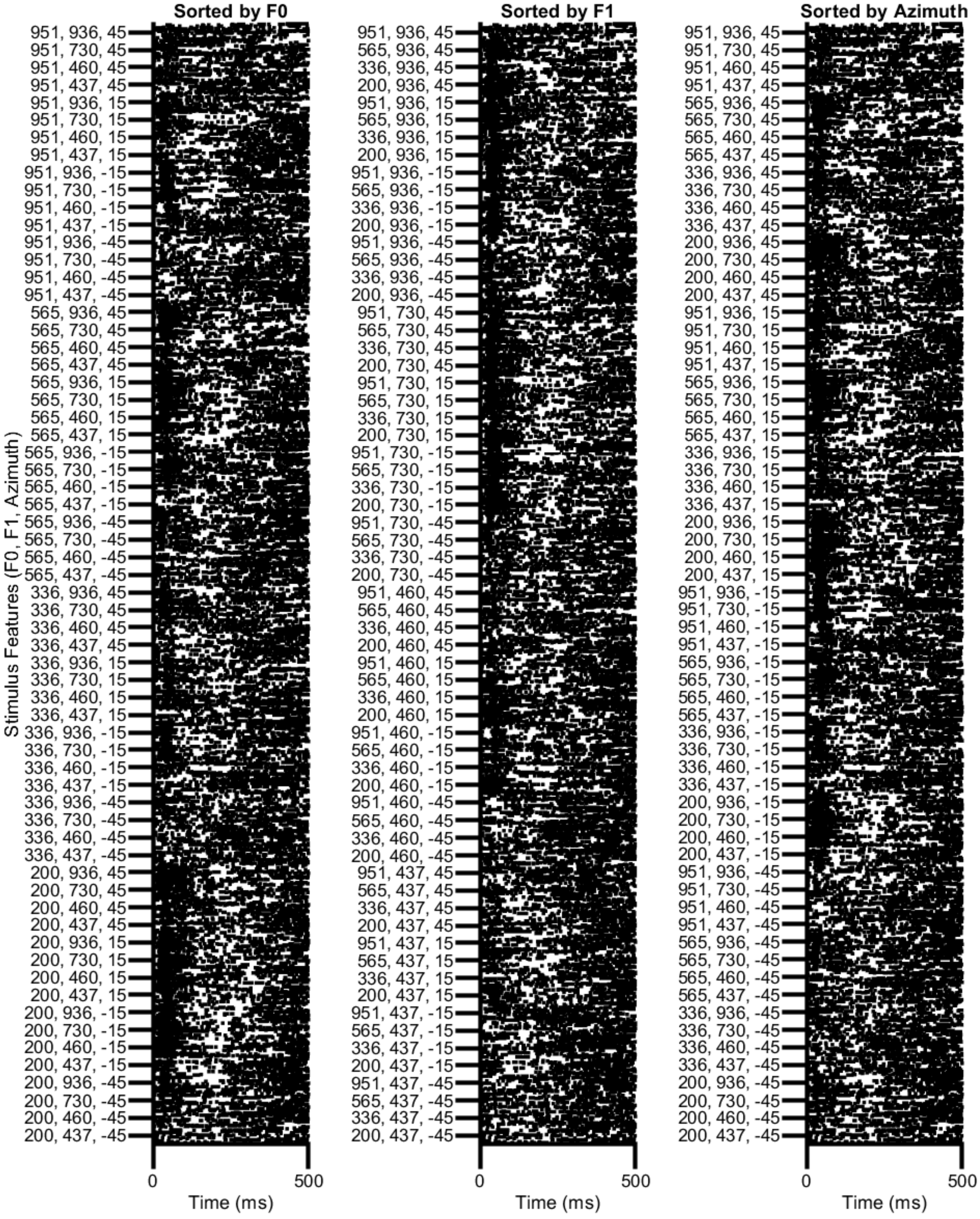

Extended data 3-2

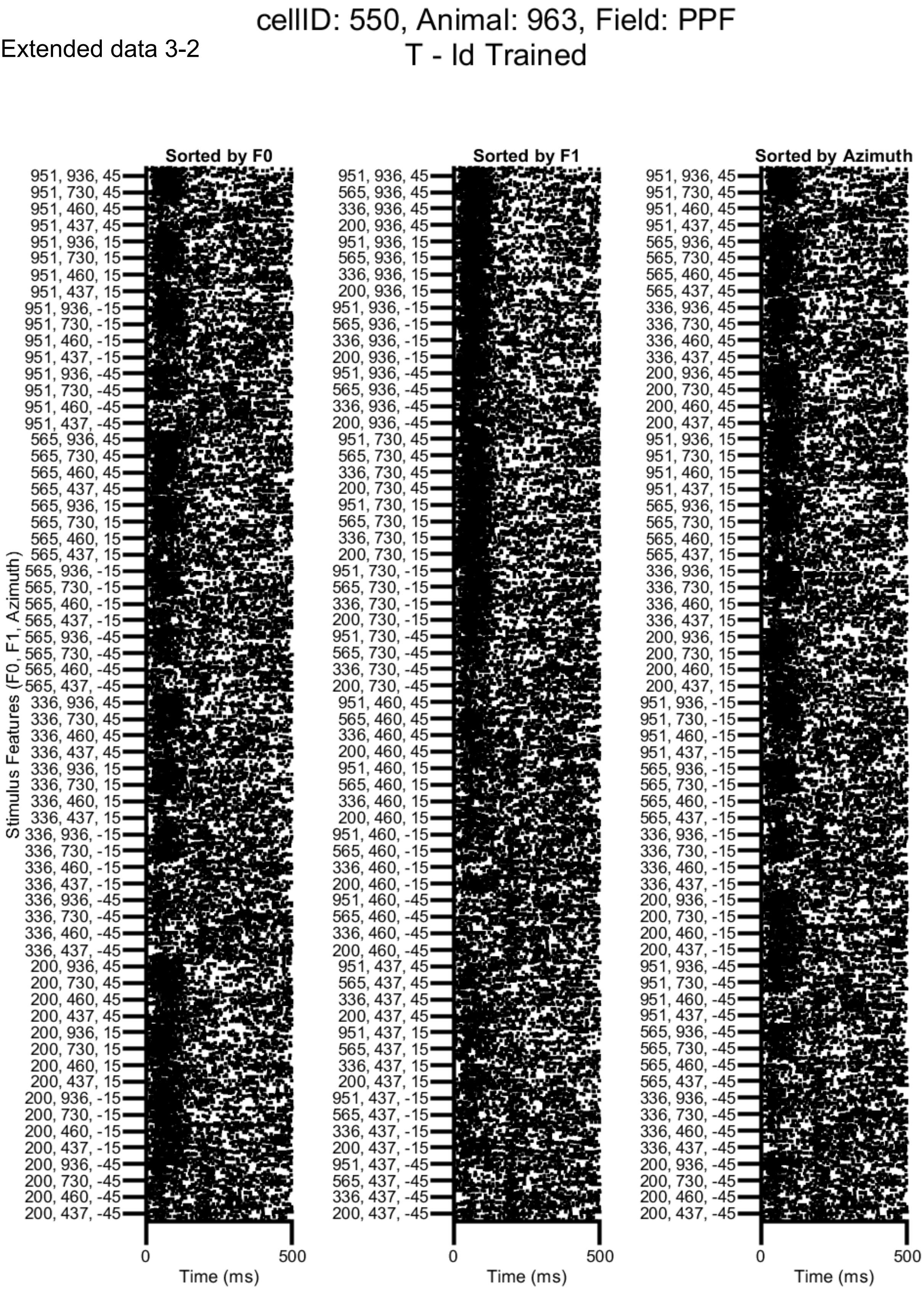

Extended data 3-3

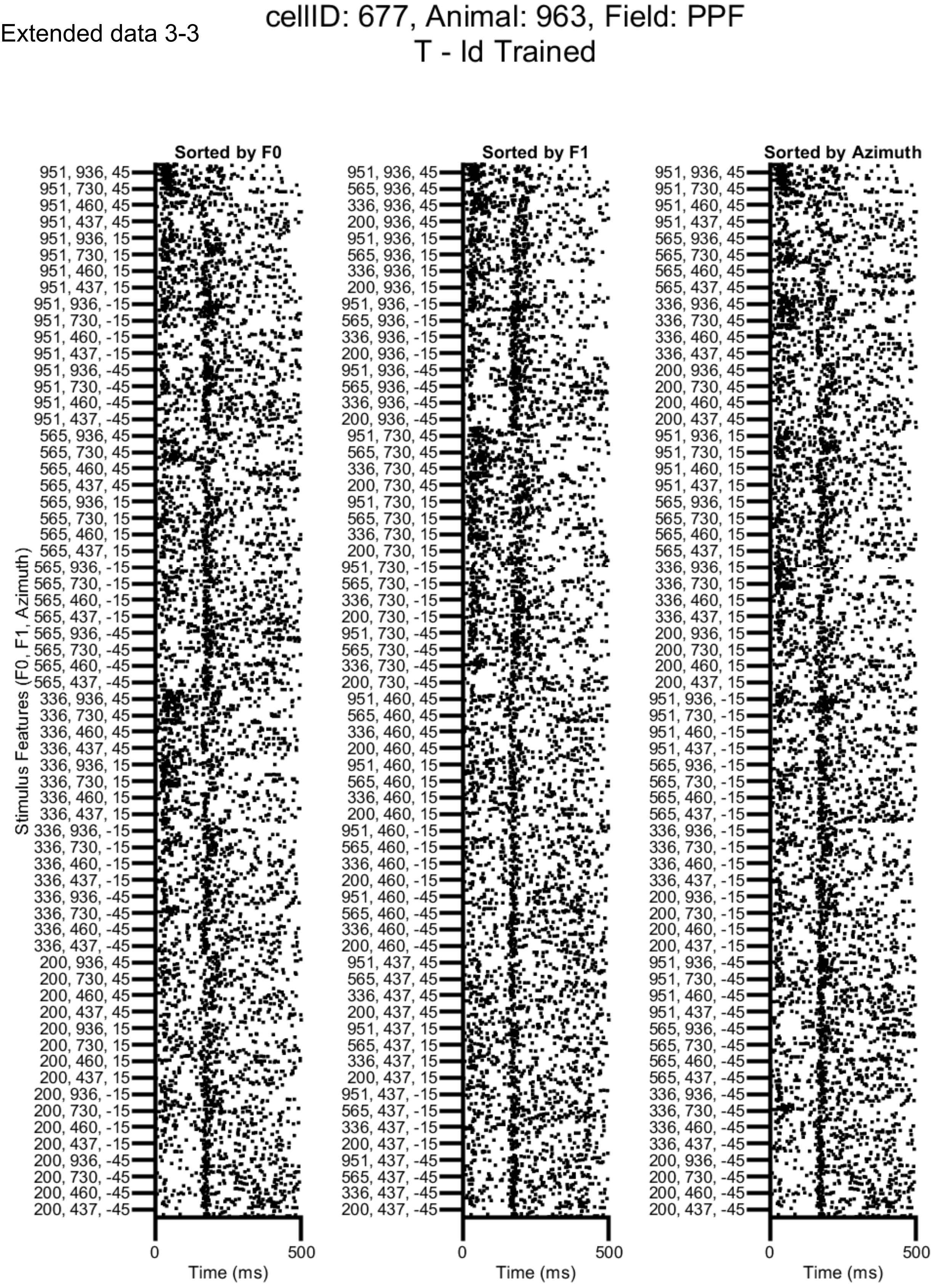

cellID: 312, Animal: 943, Field: PSF
T - ld Trained

Extended data 4-1

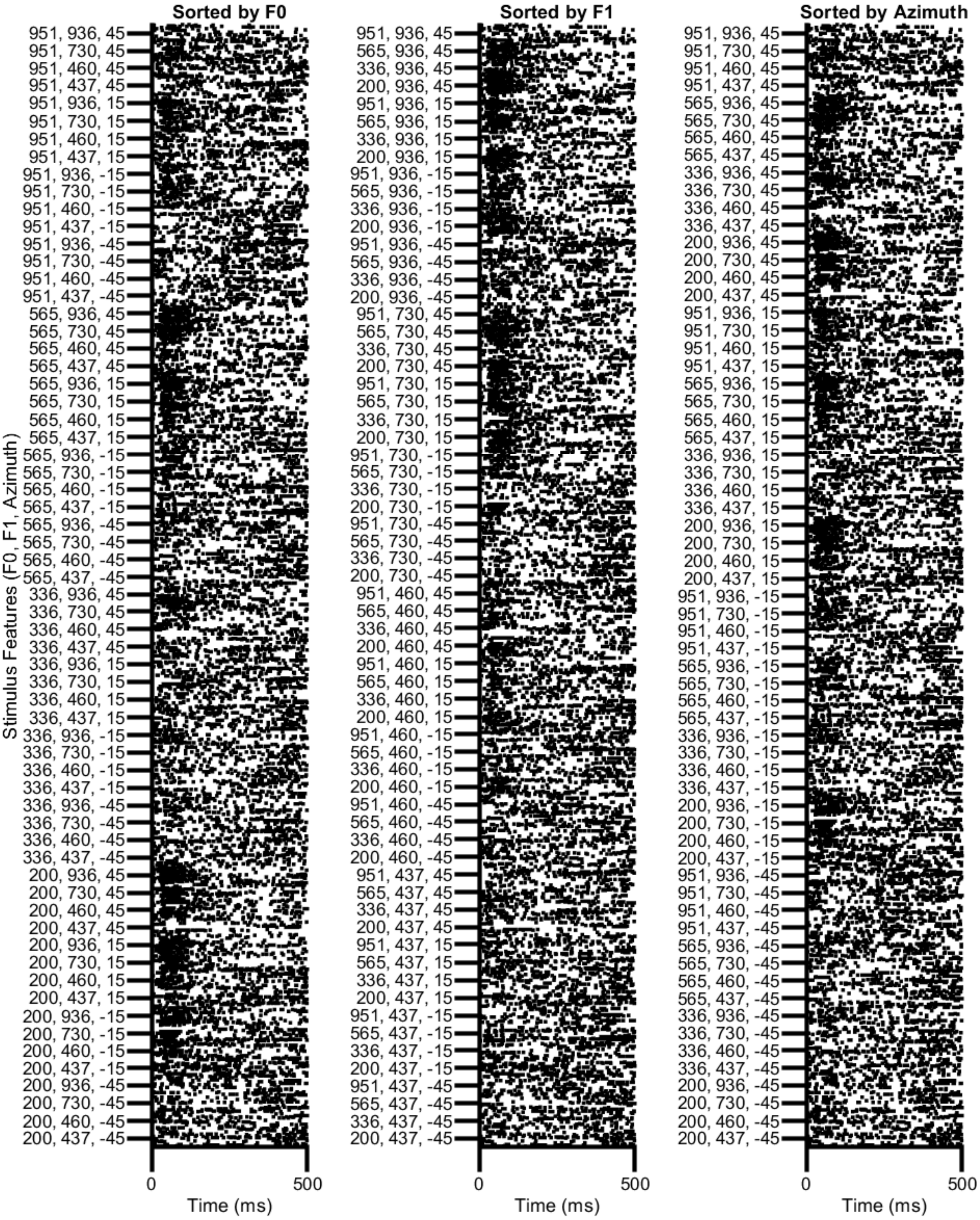

Extended data 4-2

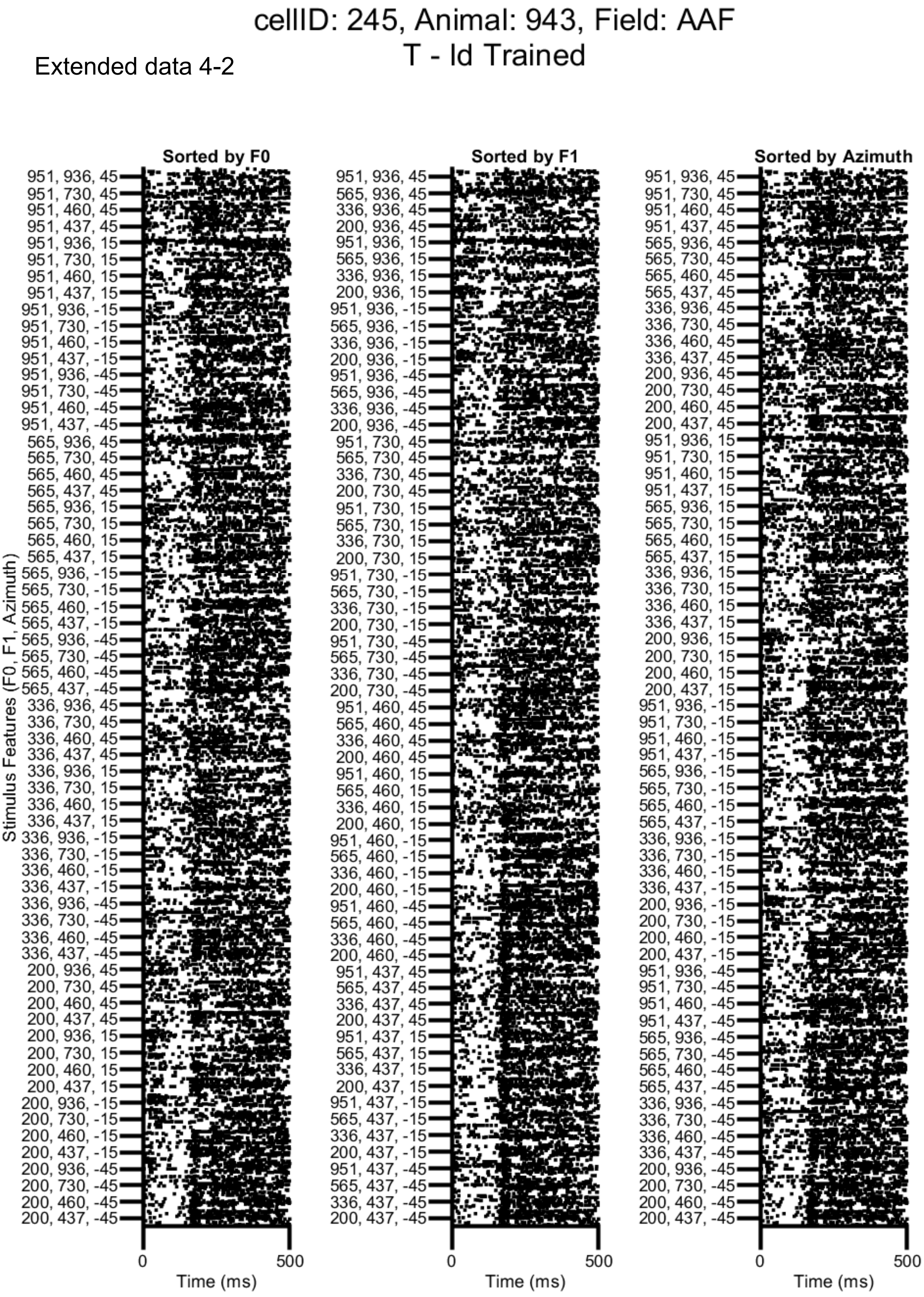


xtended data 2-2

Extended data 4-3

cellID: 538, Animal: 963, Field: PPF
T - ld Trained

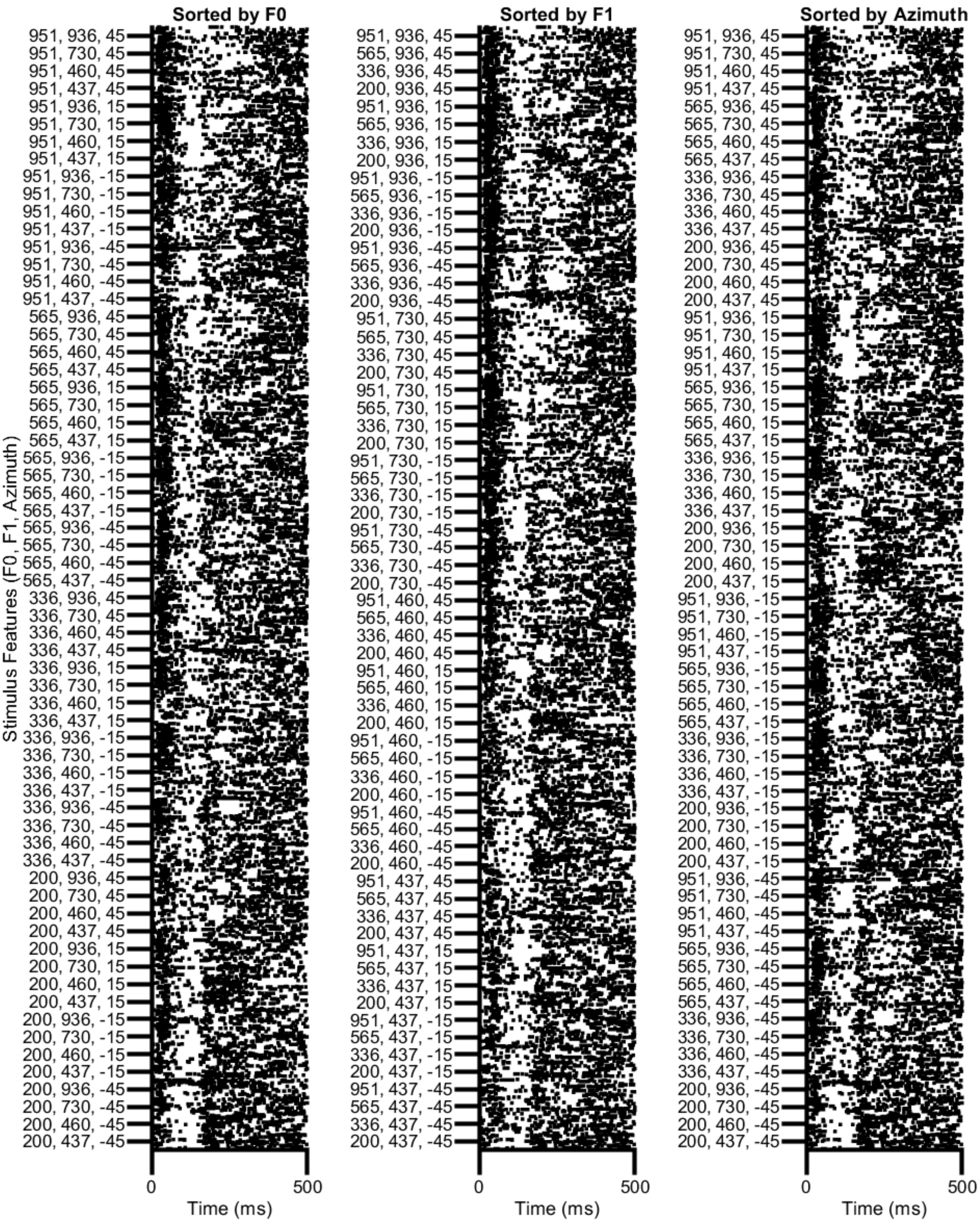

### cellID: 276, Animal: 943, Field: PSF
#### T - Id Trained

Extended data 4-4

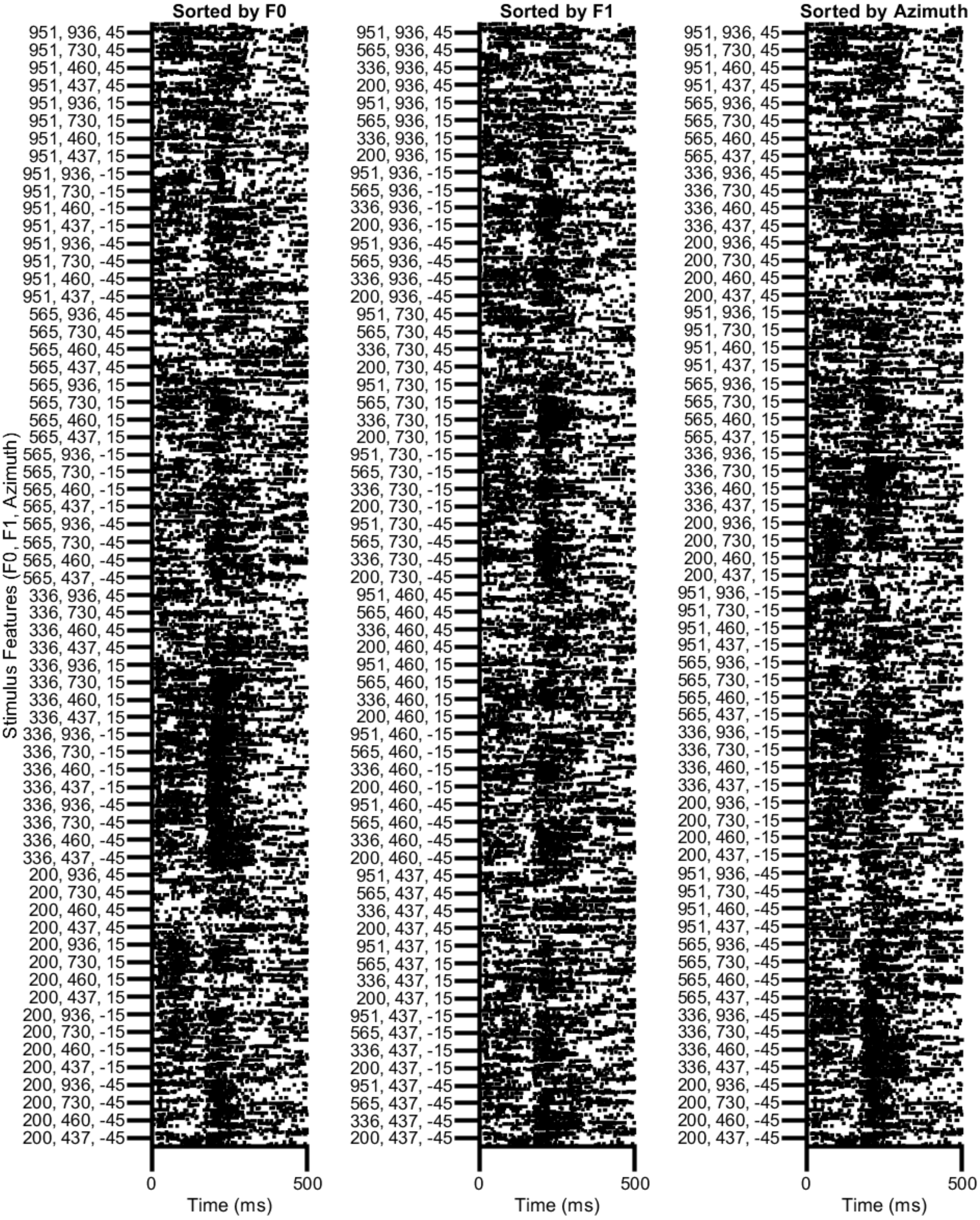

Extended data 5-1

cellID: 362, Animal: 943, Field: A1
T - ld Trained

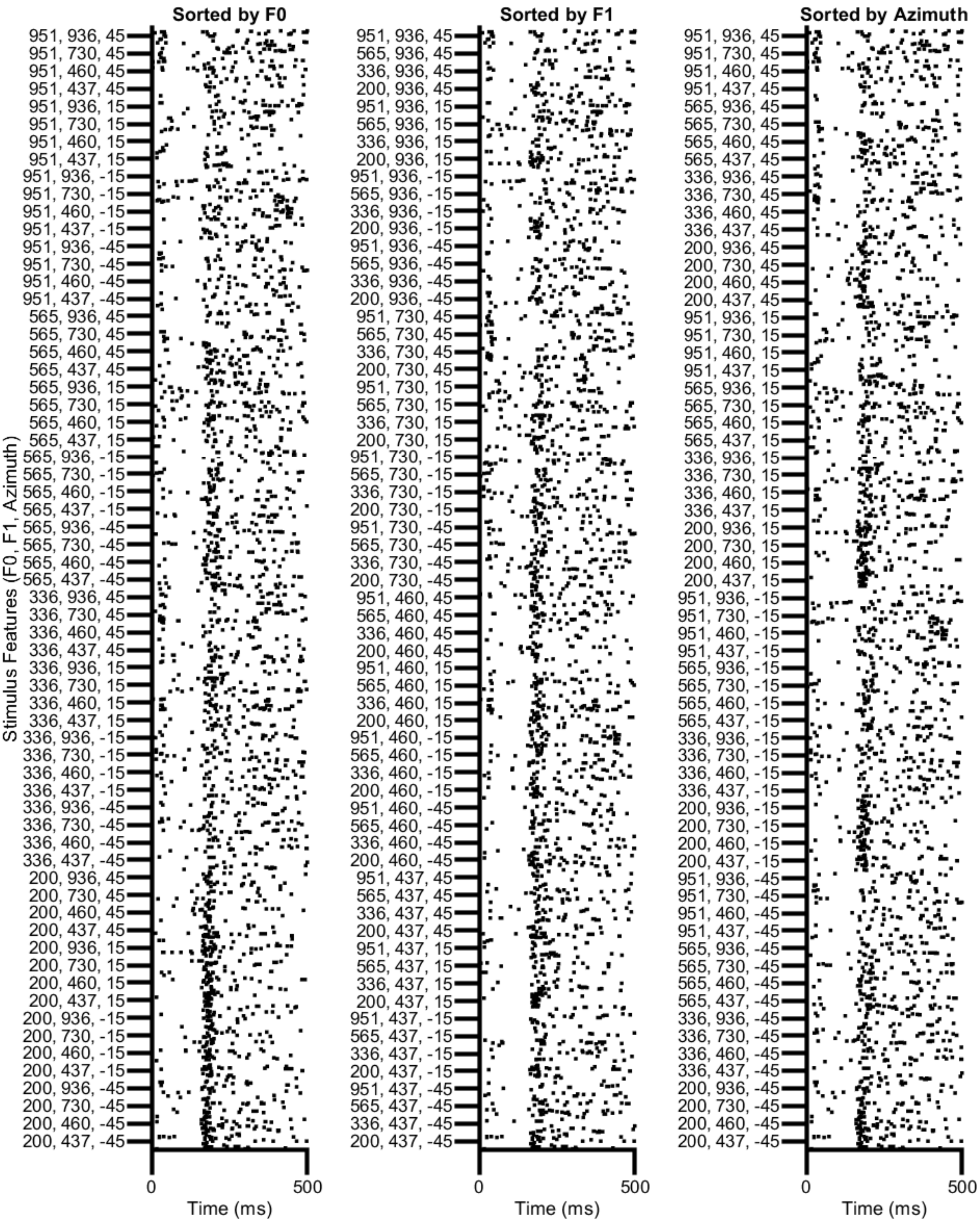

cellID: 309, Animal: 943, Field: PSF
T - Id Trained

Extended data 5-2

Extended data 6-3

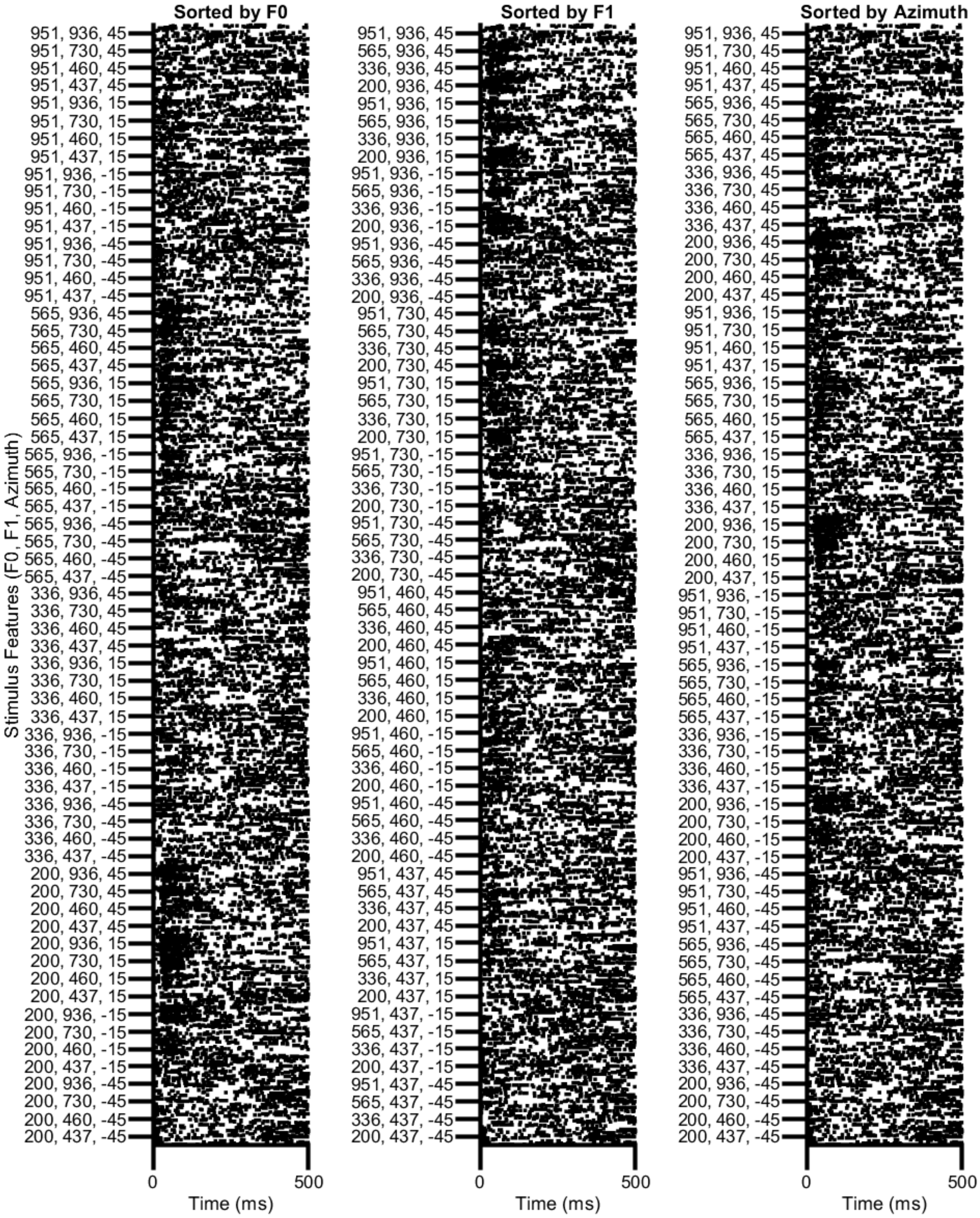


=ended data 2-2

Extended data 5-3

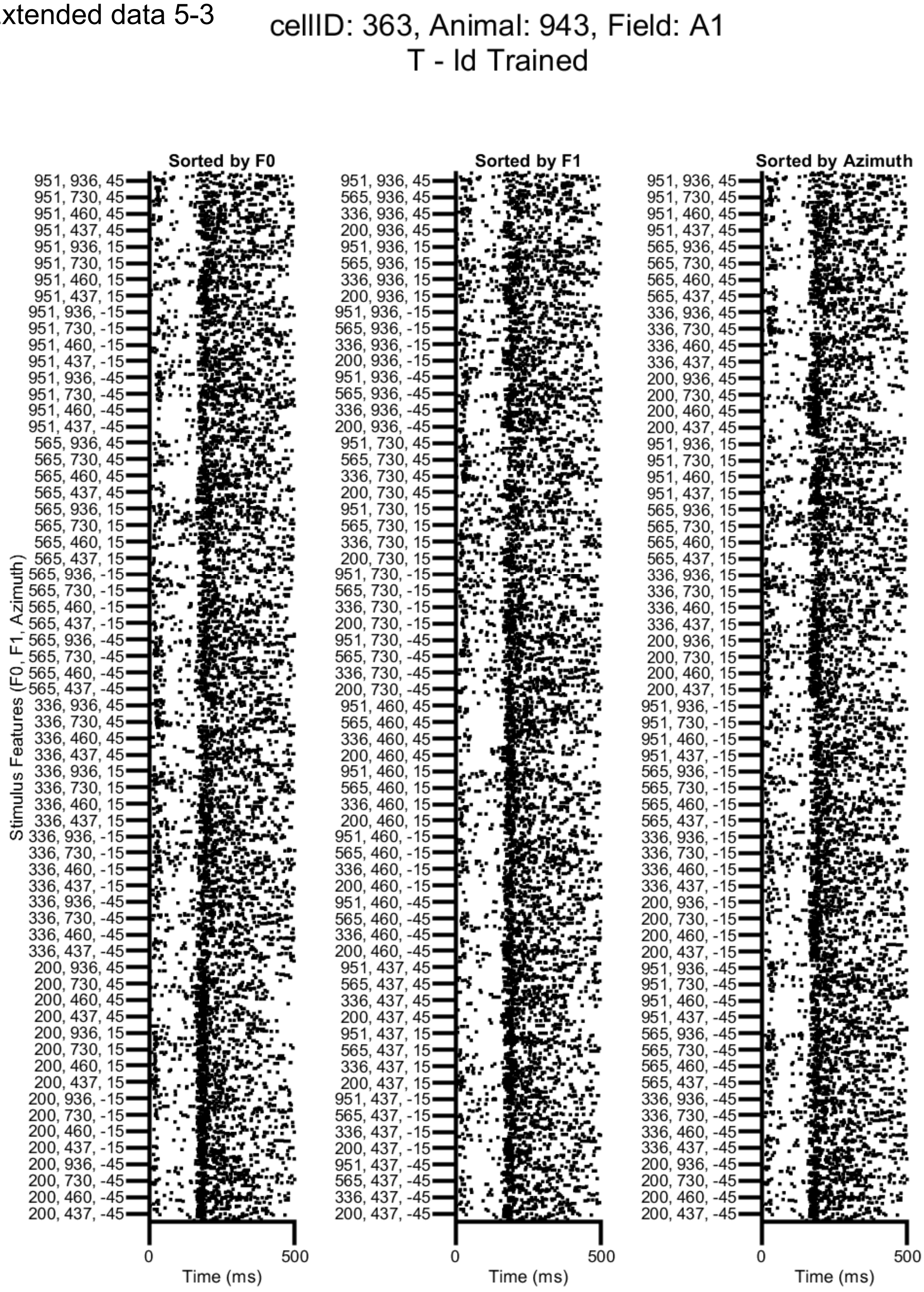

Extended data 6-1

cellID: 185, Animal: 848, Field: PSF
TP - Disc Trained

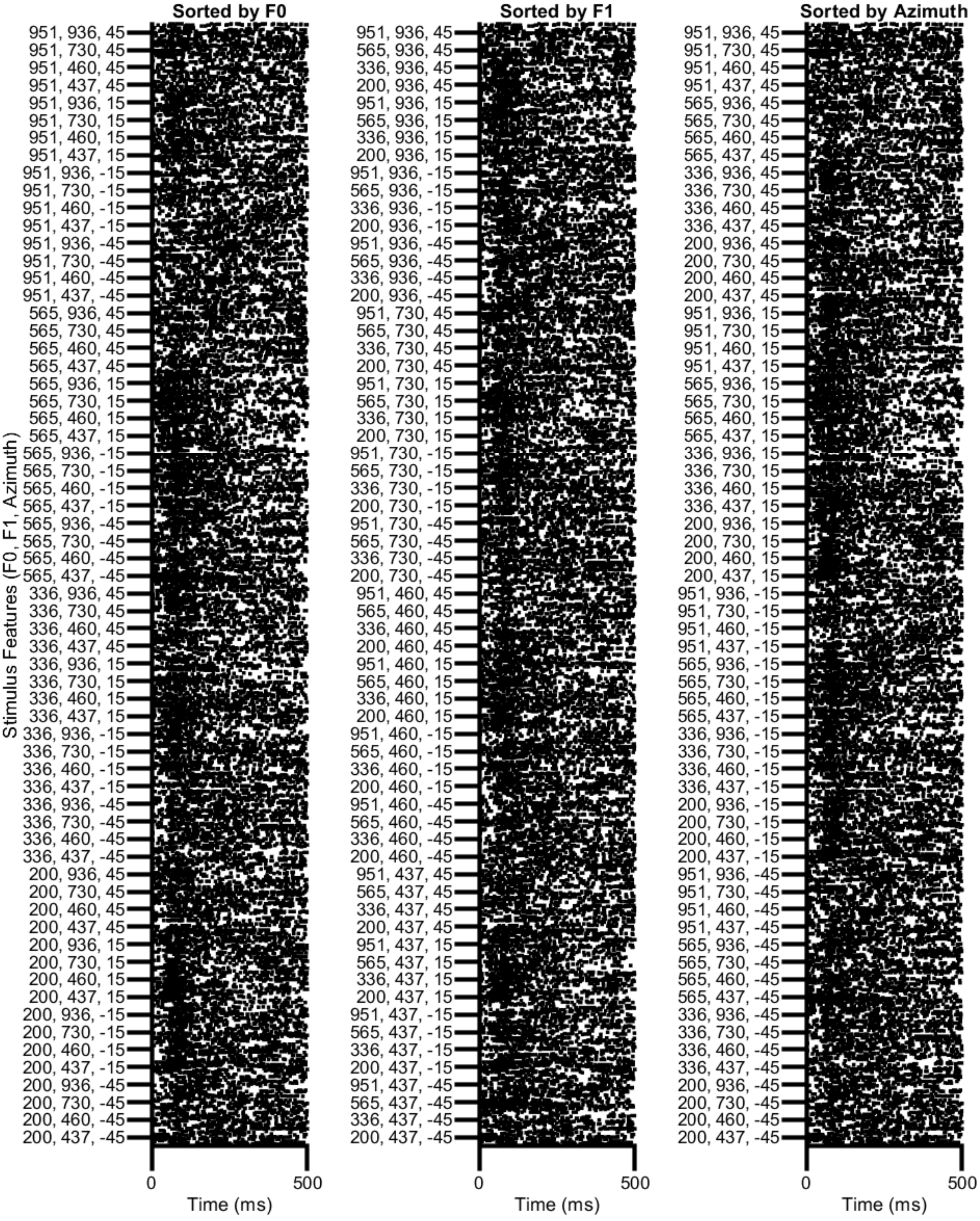

Extended data 6-2

### cellID: 82, Animal: 833, Field: PSF
#### TP - Disc Trained

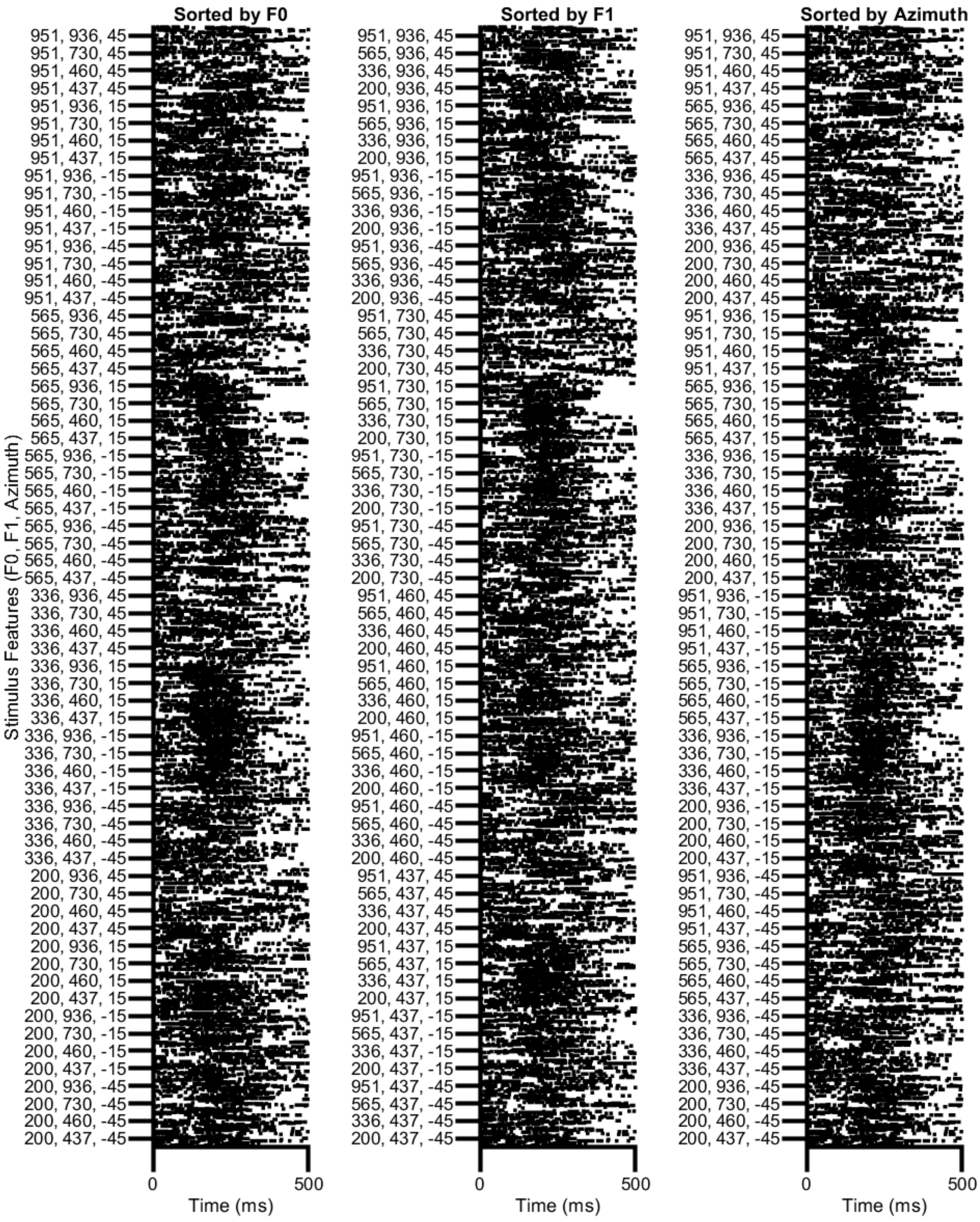
